# RNA helicase Ddx21 safeguards fetal HSPC expansion by recruiting Kdm5a to epigenetically sustain ribosomal and hematopoietic gene transcription

**DOI:** 10.1101/2025.09.04.674186

**Authors:** Adrian On-Wah Leung, Chang Li, Xianyin Huang, Zhuming Fan, Yuning Chen, Jiaxing Liu, Hong-Man Lin, Xiaoming Cui, Fujia Zhang, Ruoxi Niu, Kam-Tong Leung, Hoi-Hung Cheung

**Author notes:** These authors contributed equally to the work (Adrian On-Wah Leung, Chang Li). Correspondence: Hoi-Hung Cheung.

## Abstract

Fetal hematopoietic stem and progenitor cells (HSPCs) require active ribosome biogenesis to sustain their rapid proliferation, yet how non-structural ribosomal factors regulate this process remains unclear. Here, we identify the RNA helicase Ddx21 as a critical determinant of fetal hematopoiesis through its epigenetic role in maintaining active ribosomal transcription. Conditional knockout of *Ddx21* in murine fetal hematopoietic cells resulted in severe anemia, fetal liver hypoplasia, and depletion of HSPCs, accompanied by erythroid maturation arrest. Mechanistically, Ddx21 deficiency disrupted ribosomal and hematopoietic gene expressions, impaired translational capacity, and activated p53 pathway, triggering cell cycle arrest and apoptosis. While p53 inhibition partially rescued proliferation defects, it failed to restore rRNA transcription, pointing to p53-independent mechanisms. Multi-omics profiling revealed that Ddx21 interacts with the histone demethylase Kdm5a, co-occupying active promoters marked by H3K4me3. Loss of Ddx21 diminished H3K4me3 levels at ribosomal DNA (rDNA) and hematopoietic genes such as *cKit* and *Gata1*, downregulating their transcription. Strikingly, Kdm5a inhibition restored rRNA expression and protein translation in Ddx21-deficient cells, and combined inhibition of Kdm5a and p53 cooperatively rescued HSPC function. These findings demonstrate a paradigm wherein Ddx21 couples ribosome biogenesis to epigenetic regulation by sequestering Kdm5a at active chromatin, thereby preserving transcriptional output essential for fetal hematopoietic expansion.

## INTRODUCTION

Hematopoiesis is a dynamic process essential for lifelong production of different blood cells. During fetal development, hematopoietic stem and progenitor cells (HSPCs) undergo rapid expansion in the fetal liver (FL) to meet the escalating demand for blood cells (Popescu et al., 2019; Weissman and Shizuru, 2008). This phase relies heavily on robust protein synthesis driven by ribosome biogenesis, a tightly coordinated process involving rRNA transcription, ribosomal protein (RP) synthesis, and ribosome assembly (Ni and Buszczak, 2023). Perturbations to ribosome biogenesis disrupt cellular homeostasis, leading to ribosomopathies such as Diamond-Blackfan anemia (DBA) characterized by hematopoietic failure and congenital anomalies (Da Costa et al., 2020; Iskander et al., 2024; Ulirsch et al., 2018). While mutations in structural RP genes (e.g., *RPS19* and *RPL5*) are well-documented in ribosomopathies, the roles of non-structural ribosomal factors in regulating ribosome dynamics and their impact on hematopoiesis are underexplored. As ribosome biogenesis influences stem cell fate decision and balances self-renewal and differentiation (Lv et al., 2021; Magee and Signer, 2021; Saba et al., 2021; Zheng et al., 2024), the loss of ribosome biogenesis factors is thought to disturb the hematopoietic system. For instance, germline mutations of *DDX41*, a ribosome assemby factor for processing snoRNA, cause ineffective hematopoiesis and myelodysplasia (Bi et al., 2025; Chlon et al., 2021). However, some ribosome biogenesis factors (e.g. *HectD1*) are more tolerated when deleted in the fetal stage and display abnormality in adult HSPCs. These studies suggest a diverse role of ribosome biogenesis factors in the control of the hematopoietic system.

DDX21, a DEAD-box RNA helicase, is a key ribosome biogenesis factor implicated in rDNA transcription, rRNA processing, and ribosomal subunit maturation (Calo et al., 2015; Henning et al., 2003; Sloan et al., 2015). Knockdown of Ddx21 in preimplantation mouse embryo results in developmental aplasia (Bora et al., 2021). Moreover, DDX21 mislocalization and impaired ribosome biogenesis are observed in neural crest cells in disease models of ribosomopathies, such as Treacher Collins syndrome and DBA (Calo et al., 2018). In zebrafish, *ddx21* is also required for Vegfc-driven developmental lymphangiogenesis by balancing endothelial cell ribosome biogenesis and p53 function (Koltowska et al., 2021). These previous studies suggest a tissue-specific role of DDX21 in development. However, the function of DDX21 in fetal HSPCs, cells with extraordinary biosynthetic demands, remains undefined, nor has the role of DDX21 in mammalian hematopoiesis been elucidated.

Here, we investigate the loss-of-function of *Ddx21* during fetal hematopoiesis. Using conditional knockout (cKO) mice, we demonstrate that deletion of *Ddx21* in mouse hematopoietic cells causes severe anemia, liver hypoplasia, and HSPC depletion in the fetus. Multi-omics analyses reveal that Ddx21 interacts with the histone demethylase Kdm5a to preserve an active chromatin state at ribosomal and hematopoietic genes, thus maintaining an active transcriptional output. Our findings position DDX21 as a nexus linking ribosome biogenesis to epigenetic regulation, offering insights into the pathogenesis of ribosomopathies and highlighting therapeutic potential by targeting the ribosome-epigenome interface.

## RESULTS

### Knockout of Ddx21 in hematopoietic cells results in defective hematopoiesis and fetal anemia

As germline deletion of *Ddx21* in mice is embryonically lethal, we generated floxed *Ddx21* mice (*Ddx21^f/f^*) for cKO (Figure S1A). By crossing with *Vav1-Cre* mice, *Vav1-Cre;Ddx21^f/f^* (*Ddx21*^cKO^) showed complete deletion of *Ddx21* in fetal hematopoietic cells (Figures S1B and S1C). Notably, the proportion of viable *Ddx21*^cKO^ embryos declined dramatically after E16.5 (Figure 1A). The *Ddx21*^cKO^ embryos were pale; and the FL size was significantly smaller than the wild-type (WT) counterpart of the same stage (Figures 1B and S1D). Increased cell death was noted in the FL starting from E12.5, concomitant with the reduced cellularity (Figures 1C and 1D). Hemoglobin content per FL was also markedly decreased, indicating fetal anemia (Figure 1E). To investigate whether hematopoiesis was disrupted, we used flow cytometry (please refer to Figure S2 for gating strategy) to analyze lineage-depleted (Lin^−^) cells that contain Lin^−^Sca1^+^cKit^+^ (LSK) HSPCs and the committed Lin^−^Sca1^−^cKit^+^ (LS^−^K) progenitors. In *Ddx21*^cKO^, both LSK and LS^−^K cells, represented as percentage relative to the total FL cells, were found dramatically decreased at E13.5, and were almost undetectable at E18.5, a stark contrast to the rapid expansion in *Ddx21*^WT^ (Figure 1F). Further analysis of the LSK subpopulations including hematopoietic stem cells (HSC), multipotent progenitors MPP1 (balanced), MPP2 (megakaryocyte/erythrocyte-biased), and MPP3&4 (myelocyte/lymphocyte-biased), defined by SLAM antigens CD150/CD48 (Kiel et al., 2005), showed a marked decrease in both E13.5 and E16.5 FLs (Figure 1G). Likewise, committed LS^−^K progenitors containing CMP (common myeloid progenitor), GMP (granulocyte-monocyte progenitor) and MEP (megakaryocyte-erythroid progenitor) were also decreased (Figure 1G). Quantification of the absolute cell numbers of these stem/progenitor cells also showed significant reductions in the cKO group (Figures S3A and S3B). The depletion of HSPCs, especially erythropoietic MPP2 and MEP, likely accounted for the anemia phenotype and the decline in embryonic survival.

**Figure 1.**
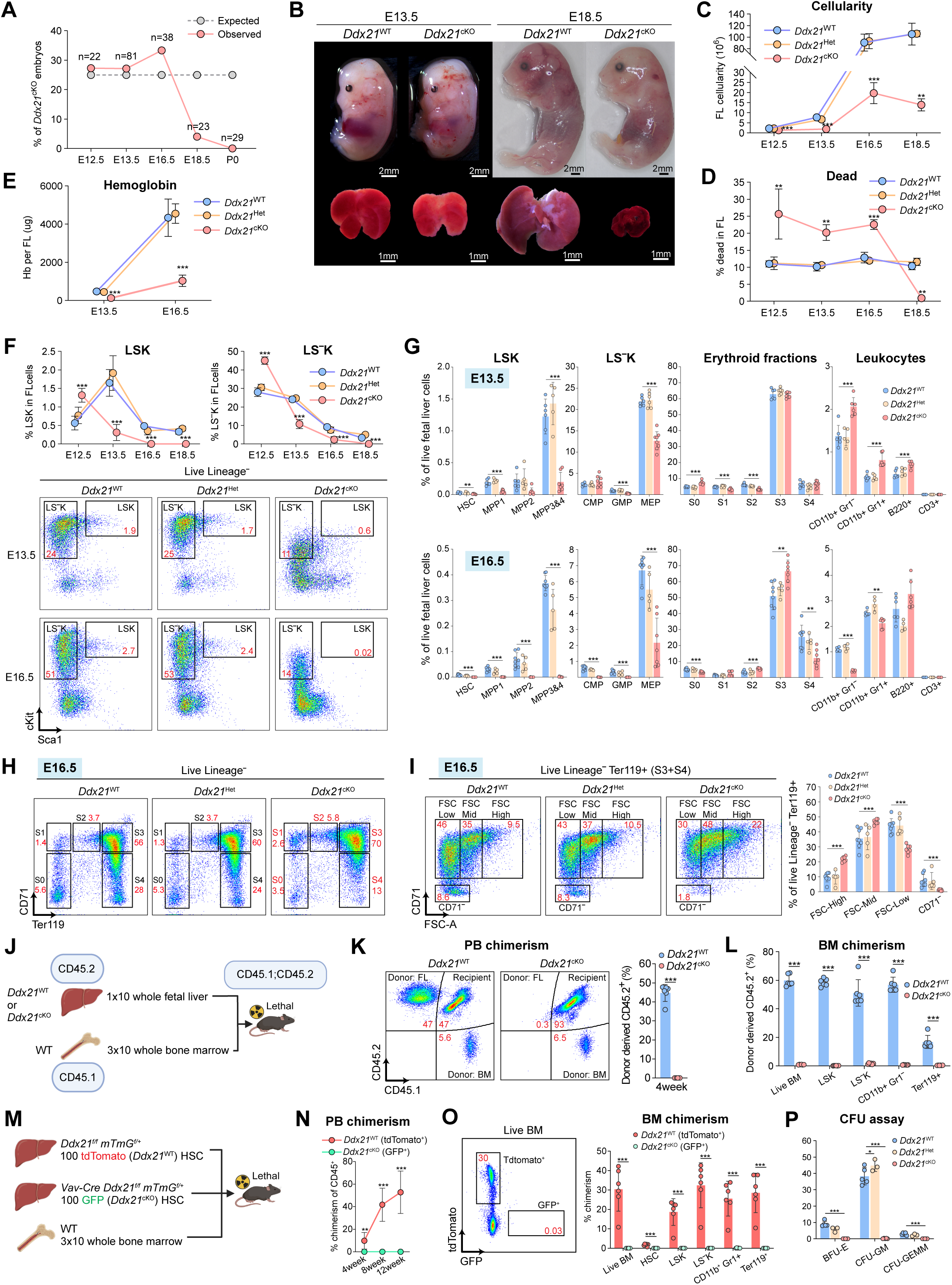
Ddx21 knockout in fetal hematopoietic cells results in severe hematopoietic defect and embryonic lethality.

### Effect of Ddx21 knockout on erythroid maturation

To further investigate the effect of Ddx21 loss on erythropoiesis (the predominant hematopoietic event in the fetus), we analyzed the maturation of FL erythroblasts by gating the Lin^−^ cells and analyzed the erythroid fractions (S0-S4) using the surface markers CD71/Ter119 (Figure S2) (Chen et al., 2009). At E16.5, the absolute and relative numbers of early committed erythroblasts (S1, Ter119^−^CD71^high^) and late immature erythroblasts (S2 & S3, Ter119^+^CD71^high^) were increased in *Ddx21*^cKO^, concomitant with the decrease in mature erythrocytes (S4, Ter119^+^CD71^low^) (Figures 1G, 1H and S3B). Among the late erythroid populations (S3+S4), FSC-based cell size analysis provides a higher resolution of the terminal differentiation where a shift towards the less mature and larger sized erythroblast (FSC^high^) was observed in *Ddx21*^cKO^. As a result, terminally differentiated reticulocytes/erythrocytes (Ter119^+^CD71^−^) were decreased (Figures 1I and S3C). These observations suggest a maturation delay or differentiation blockage during terminal erythropoiesis.

### Knockout of Ddx21 abolishes hematopoietic reconstitution

The FL contains definitive HSPCs that can reconstitute the bone marrow (BM) and differentiate into myeloid and lymphoid lineages upon transplantation to myeloablated recipient mice, a golden standard to assess the hematopoietic capability of HSPCs (Rebel et al., 1996). We performed competitive bone marrow transplantation (BMT) by co-injecting 1.5×10^6^ live FL cells from *Ddx21*^WT^ or *Ddx21*^cKO^ E13.5 embryos (CD45.2) with 3×10^5^ competitive WT BM cells (CD45.1) into lethally irradiated adult recipients (CD45.1;CD45.2) (Figures 1J and S4A). As anticipated, we observed a high peripheral blood (PB) chimerism in mice injected with *Ddx21*^WT^ one month post-transplantation. However, PB chimerism for *Ddx21*^cKO^ was less than 1% (Figure 1K). Similarly, chimerism in BM was significantly low for Ddx21^cKO^ – the HSPC fractions (LSK and LS^−^K) were virtually undetectable, indicating a failure in engrafting and reconstituting the BM (Figure 1L).

To address the difference in the actual HSPC numbers between *Ddx21*^WT^ and *Ddx21*^cKO^ FL that may mask the transplantation results, a competitive co-transplantation experiment using equal number of HSCs was performed. We transplanted 100 tdTomato^+^ live HSCs (LSK CD150^+^ CD48^−^) of *Ddx21*^WT^ (*Ddx21^f/f^*;*mTmG*^f/+^) and 100 GFP^+^ HSCs of *Ddx21*^cKO^ (*Vav1-Cre*;*Ddx21^f/f^*;*mTmG^f+f^*) with 3×10^5^ BM cells (WT) to myeloablated adult mice. PB chimerism was monitored every 4 weeks for 3 months (Figures 1M and S4B). Similar to FL transplantation, chimerism contributed by *Ddx21*^cKO^ HSCs was less than 0.1% in both PB and BM 3 months post-transplantation, indicating an essential role of Ddx21 in HSC’s reconstitution capacity (Figures 1N and 1O). The loss of hematopoietic function in *Ddx21*^cKO^ was further supported by colony-forming unit (CFU) assay which showed a failure in generating any types of the analyzed colonies: BFU-E, CFU-GM, and CFU-GEMM, using E13.5 FL cells (Figures 1P and S4C). Together, these results suggest a cell-autonomous function of Ddx21 in safeguarding HPSCs by potentially regulating HSPC expansion during fetal hematopoiesis.

### Activation of p53 and apoptosis in Ddx21-defecient HSPCs

DDX21 loss is known to induce p53-dependent cell cycle arrest in mutant zebrafish and cultured cancer cells (Koltowska et al., 2021; McRae et al., 2020), providing a rationale accounting for our observed hematopoietic failure. However, as these molecular events were reported to be cell type- and species-specific (Calo et al., 2018), the regulatory axis in HSPCs remained elusive. We thus performed RNA-seq and analyzed the transcriptome of Lin^−^cKit^+^ (LK) HSPCs, and observed a pronounced upregulation of p53 downstream genes such as *Cdkn1a*, *Fas* and *Mdm2* in *Ddx21*^cKO^ (Figure 2A). Both Kyoto Encyclopedia of Genes and Genomes (KEGG) and Gene Set Enrichment Analysis (GSEA) revealed the enrichment of p53 signaling pathway upon *Ddx21* knockout (Figures 2B and 2C). Further analysis of the p53 downstream targets revealed a concerted upregulation of apoptotic signaling genes and inhibition of cell cycle progression genes (Figure 2D).

**Figure 2.**
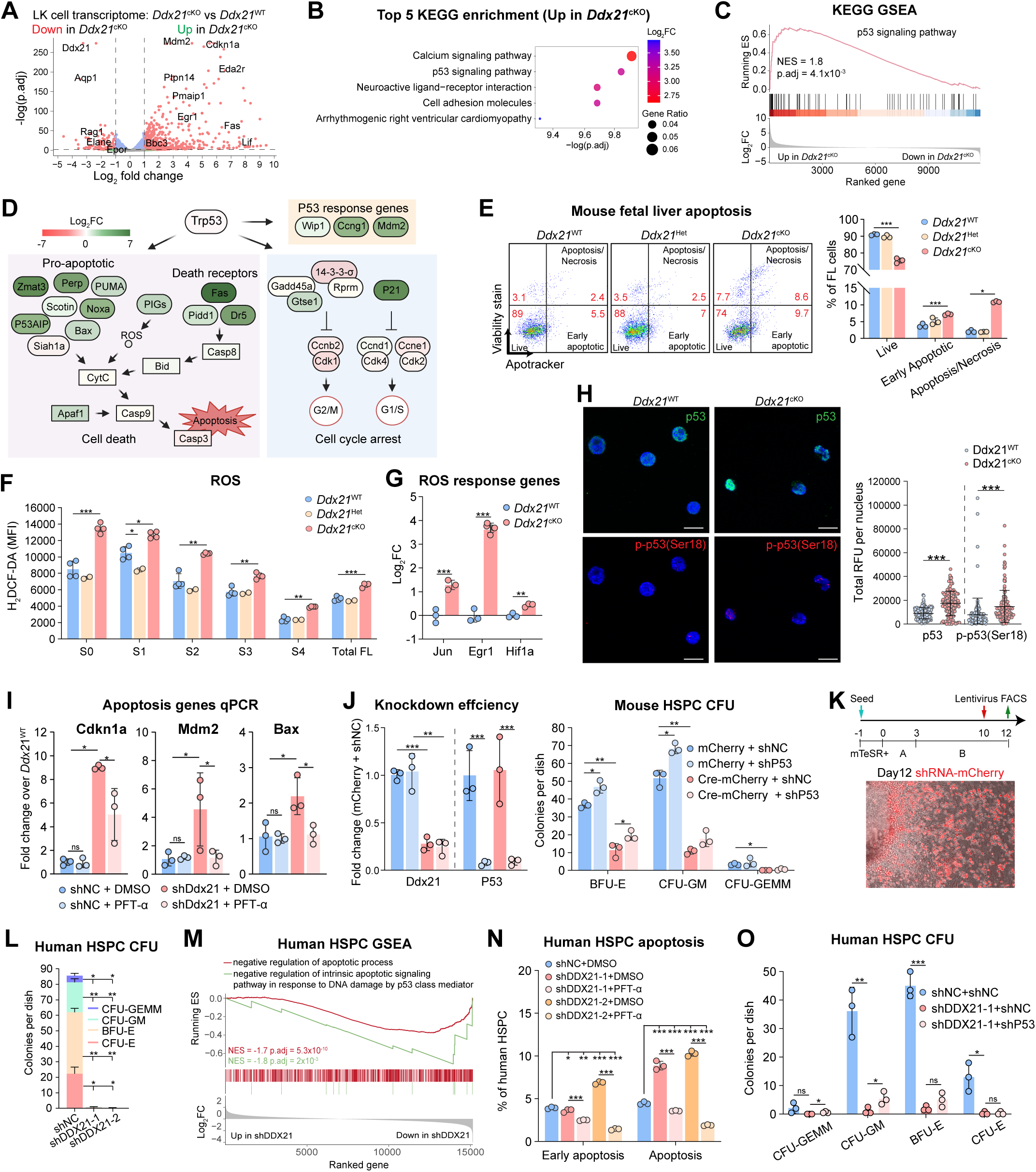
p53 inhibition prevents ROS and p-p53(Ser18)-driven apoptosis and partially restores hematopoietic function.

To examine the apoptotic changes, E13.5 FL cells were analyzed by flow cytometry using Apotracker (an Annexin-V equivalent dye) and dead cell probe FVS (fixable viability stain). In line with the transcriptome, a significant increase in early apoptosis (Apotracker^+^ FVS^−^) and late apoptosis/necrosis (Apotracker^+^ FVS^+^) was observed (Figure 2E). Studies have demonstrated that high level of reactive oxygen species (ROS) triggers p53 phosphorylation at Ser18 (Ser15 for human P53), a molecular marker of p53 activation (Assaily et al., 2011). We analyzed the ROS level in sorted live cells from E16.5 *Ddx21*^cKO^ FL. The ROS level was overall elevated, especially in the erythroid progenitors (Figure 2F). Consistently, ROS response genes such as *Jun*, *Egr1* and *Hif1a* were found upregulated (Figure 2G). Additionally, immunofluorescent staining of the p53 protein revealed elevated nuclear signals of p53 in *Ddx21*^cKO^ LK cells, and a similar change was observed for the p-p53(Ser18) (Figure 2H).

### Inhibition of p53 alleviates cell death but not the hematopoietic function

To evaluate the role of p53 in mediating the loss of Ddx21, we inhibited p53 by Pifithrin-α (PFT-α), and successfully suppressed p53 downstream genes such as *Cdkn1a*/p21, *Mdm2* and *Bax* in Ddx21 knockdown HSPC (Figure 2I). Next, we tested whether p53 inhibition in *Ddx21*-deleted HSPCs (mediated by p53-shRNA and Cre-mCherry lentiviruses) was sufficient to rescue the hematopoietic defect. Despite a high efficiency of *p53* knockdown in LK cells, CFU assay showed a partial rescue for the BFU-E and CFU-GM colonies (Figure 2J).

To recapitulate the findings in human hematopoietic cells, we differentiated human HSPCs (CD45^+^CD34^+^CD43^+^) from pluripotent stem cells and silenced *DDX21* by two different shRNAs (shDDX21-1 and shDDX21-2) (Figures 2K, S5A and S5B). Consistent with mouse *Ddx21* knockout, *DDX21* knockdowns in human HSPCs failed to form hematopoietic colonies (Figures 2L). GSEA analysis comparing *DDX21* knockdown and non-targeting control (shNC) transcriptomes also highlighted the activation of apoptosis (Figure 2M). In line with mouse p53 inhibition, PFT-α prevented apoptosis in *DDX21* knockdown HSPC (Figure 2N). Similarly, p53 knockdown in human HSPCs could only slightly restore hematopoietic capabilities as demonstrated by CFU assay (Figure 2O). These results indicate that p53-dependent apoptosis is not the major cause for hematopoietic failure in DDX21-deficient cells.

### Ddx21 loss downregulates ribosome biogenesis in HSPCs

Ddx21 is a key non-structural ribosome biogenesis factor implicated in rDNA transcription (Calo et al., 2015; Xing et al., 2017). To examine the impact of Ddx21 loss in ribosome biosynthesis, GSEA analysis of the mouse HSPC transcriptome revealed enrichment of “ribosome biogenesis” gene ontology (GO) term (Figure 3A). Among the significantly differentially expressed genes (DEGs) (p.adj < 0.05) in the ribosome biogenesis GO term, most genes were downregulated in *Ddx21*^cKO^ (Figure 3B). We also analyzed the same gene set for human HSPC, and revealed 43 overlapping DEGs between the two species (Figures 3C and S5C). As “structural constituent of ribosome” GO term was also enriched (Figure 3A), we analyzed the genes encoding RP and rRNA. In *Ddx21*^cKO^ HSPCs, approximately one-third of all expressed RP genes were downregulated (Figure 3D). Consistently, rRNA expression (pre-rRNA, 18S and 28S rRNA) was suppressed, which could not be restored by blocking p53 with PFT-α (Figures 3E, 3F and S5D).

**Figure 3.**
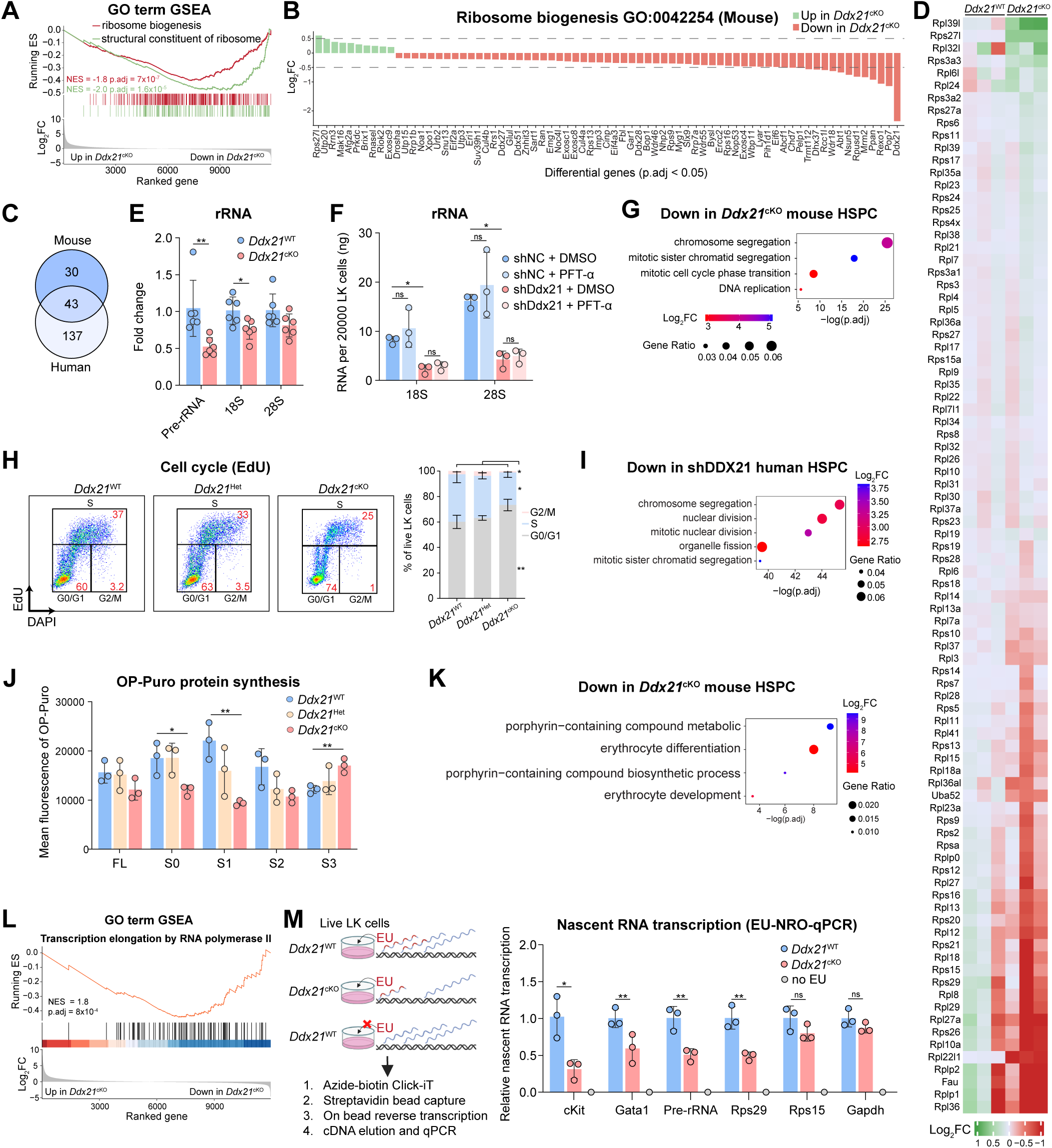
Ddx21 loss inhibits protein synthesis and RNA transcription in hematopoietic progenitors.

In addition to apoptosis, the lack of hematopoietic expansion in *Ddx21* knockout may be attributed by cell cycle arrest in the progenitors. Transcriptome analysis revealed significant enrichment of mitosis and DNA replication related GO terms (Figure 3G). To examine the changes in cell cycle and DNA replication *in vivo*, we injected EdU to the pregnant mice at E13.5 and harvested FL cells for cell cycle analysis. Both LS^−^K progenitors and erythroid progenitors (S0-S1) showed an increase in G0/G1 phase and concomintant decreases in S and G2/M phases, indicating a blockage at G1-S transition (Figures 3H and S6A). Consistently, *DDX21* knockdown in human HSPC also showed significant enrichment of mitosis-related GO terms (Figure 3I).

### Suppression of protein synthesis and transcription in Ddx21 knockout HSPCs

The downregulation of ribosome biogenesis in *Ddx21*^cKO^ HPSCs suggest a potential change in the protein translation capacity. By O-propargyl-puromycin (OPP) incorporation assay, *Ddx21*^cKO^ FL cells showed decreased protein synthesis. A significant reduction was observed in the erythroid progenitors (S0/S1) (Figure 3J). As erythroid progenitors are metabolically active in translation (Magee and Signer, 2021; Zheng et al., 2024), the reduction in protein synthesis coincided with the downregulation of porphyrin compound metabolism and erythrocyte differentiation (Figure 3K). Furthermore, we found downregulation of key hematopoietic/erythropoietic genes such as *cKit* and *Gata1*, in addition to several DBA-related genes (Figures S6C and S6D). Of note, GSEA analysis revealed negative enrichment of GO term “transcription elongation by RNA polymerase II” (Figure 3L). These observations prompted us to determine whether Ddx21 regulates RNA transcription. By 5-ethynyluridine (EU) incorporation and Click-iT chemistry, we measured nascent RNA synthesis in cultured LK cells. Nascent pre-rRNA and *Rps29* transcripts were decreased in *Ddx21*^cKO^ cells. Similarly, nascent transcripts of hematopoietic genes *cKit* and *Gata1* were also significantly decreased (Figure 3M). These results suggest that Ddx21 controls transcription of key hematopoietic factors important for HSPC proliferation and differentiation, in addition to the its generic function in ribosome biogenesis.

### Ddx21 interacts with histone modification enzyme Kdm5a

Although p53 inhibition can rescue apoptosis in Ddx21-deficient HSPCs, it failed to rescue rRNA expression and hematopoietic colony formation. The reason for this intriguing phenomenon is unclear. To dissect the molecular network in which DDX21 interacts with, we performed BioID-MS assay in K562 cells to identify proteins interacting with DDX21 (Figures 4A and S7A-C). DDX21 was found to interact with different categories of proteins involved in, consistent with previously reported, ribonucleoprotein complex biogenesis, ribosome biogenesis, rRNA metabolic process, and RNA splicing (Figure 4B) (Lessard et al., 2018; Miao et al., 2023; Sloan et al., 2015; Xing et al., 2017). Interestingly, we found a group of histone modification enzymes enriched in the DDX21 interactome, such as H3K4me3 demethylase KDM5A and histone deacetylases HDAC1/HDAC2 (Figures 4C and S7D). Knockout of *Ddx21* in mouse HSPC had a mild effect on Kdm5a protein and little effect on Hdac1 protein (Figure S9C). Co-immunoprecipitation (co-IP) experiments demonstrated the interactions of DDX21 with KDM5A, HDAC1, HDAC2, but not SIN3A (Figures 4D, 4E, S8A-D). Disruption of the ATPase/helicase activity of the DDX21 did not abolish the DDX21-KDM5A interaction, as both helicase-dead (DDX21^DEV^) and ATPase-dead (DDX21^DAT^ and DDX21^SAT^) DDX21 mutants could also co-precipitate KDM5A (Figure 4H). Additionally, native co-IP using antibodies against endogenous proteins showed that the DDX21 immunoprecipitates contained a high amount of KDM5A but less HDAC1 and SIN3A (Figures 4F and S8E). To test whether DDX21 directly binds KDM5A and HDAC1, we performed protein pulldown experiments using purified recombinant proteins. The results showed that DDX21 could directly bind KDM5A and HDAC1 (Figures 4G and S8F). Additionally, we examined the *in situ* protein-protein interactions in freshly isolated mouse LK cells by proximity ligation assay (PLA). We observed high PLA signals for Ddx21-Kdm5a but relatively weaker for Ddx21-Hdac1/2 and Ddx21-Sin3a (Figures 4J and S9B). Next, we examined the effect of removing RNA on the protein-protein interactions. RNase A treatment did not change RFU signals of Ddx21 and its interaction partners (Figures 4I and S9A). Notably, depletion of RNA by RNase A treatment decreased the Ddx21-Kdm5a but increased the Ddx21-Hdac1 PLA signals (Figure 4J). Since in normal condition Ddx21 is predominantly recruited to RNA-rich chromatin (e.g. nucleoli), the removal of chromatin RNA may disfavor Ddx21-Kdm5a interaction. In agreement with the PLA result, co-IP experiments also showed reduced DDX21-KDM5A complex formation with RNase A treatments (Figures 4D).

**Figure 4.**
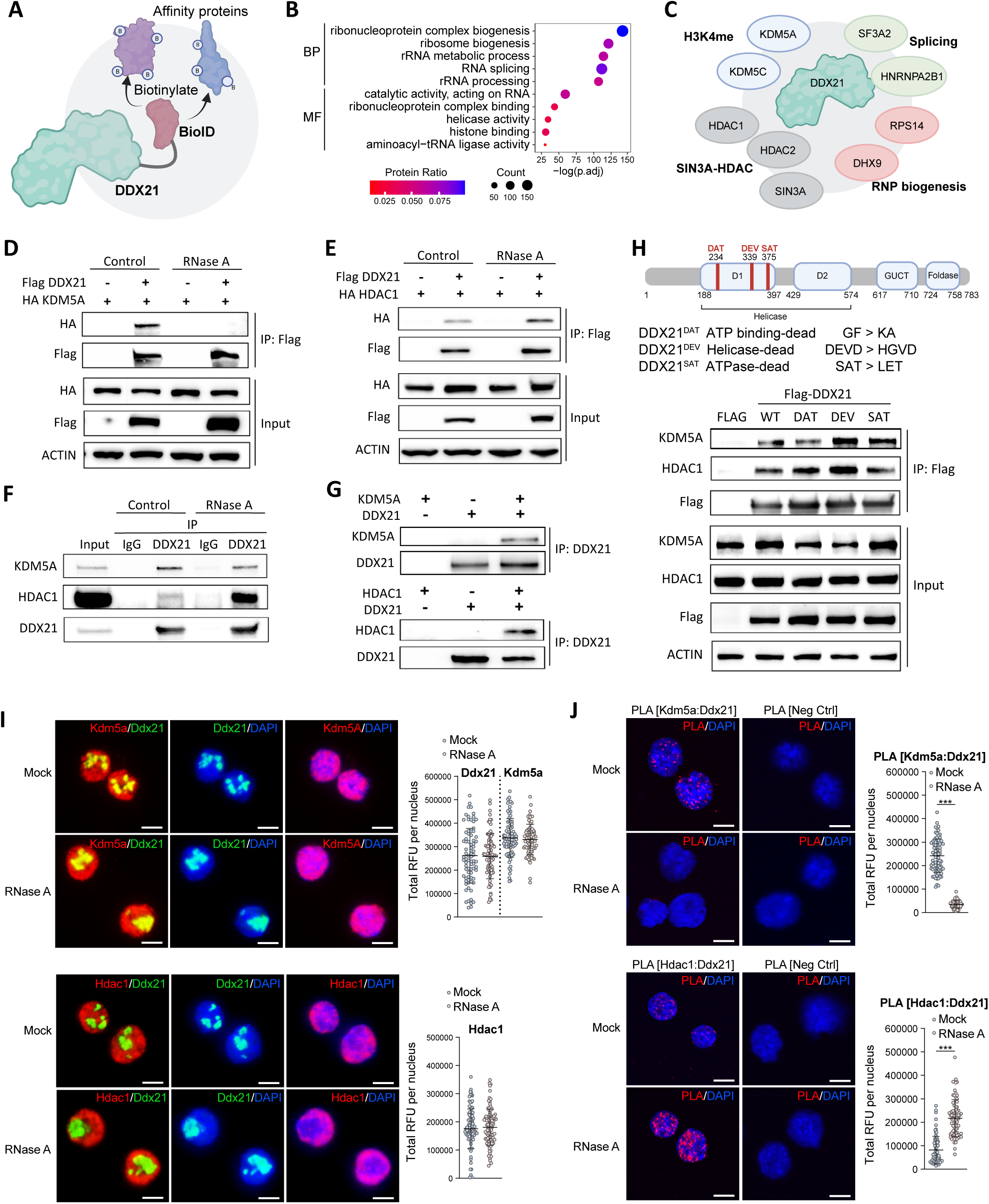
DDX21 interacts with histone demethylase Kdm5a in an RNA-dependent manner.

### Ddx21 and Kdm5a co-occupy active promoters

The above results suggest that on active chromatin where RNA is abundant, DDX21 is recruited together with KDM5A. DDX21 may sequester KDM5A from accessing to H3K4me3, preventing KDM5A-mediated demethylation and thus preserving an active chromatin state to favor transcription. To test this hypothesis, we performed CUT&Tag to map the chromatin binding sites of Ddx21 and Kdm5a, and the associated epigenetic marks for active (H3K4me3), repressive (H3K27me3) and enhancer-associated (H3K27ac, H3K4me1) histone marks in fetal HSPC. As expected, global Kdm5a binding is highly correlated with open (H3K4me3 and ATAC) but not repressive chromatins (H3K27me3) (Figure 5A), in line with a reported role of KDM5A in transcriptional regulation (Benevolenskaya et al., 2005; Brier et al., 2017). Interestingly, Ddx21-bound peaks highly overlapped with Kdm5a binding (Figure 5B). Peaks co-bound by Ddx21 and Kdm5a were mainly active (H3K4me3) promoter (Figure 5C). The promoter-associated Kdm5a binding peaks were highly correlated with Ddx21 (*R* = 0.85, based on peak intensity), H3K4me3 (*R* = 0.91) and RNA level (*R* = 0.77) (Figure 5D). These analyses suggest that Ddx21 and Kdm5a co-occupy active promoters and transcribed genes (Figure S10).

**Figure 5.**
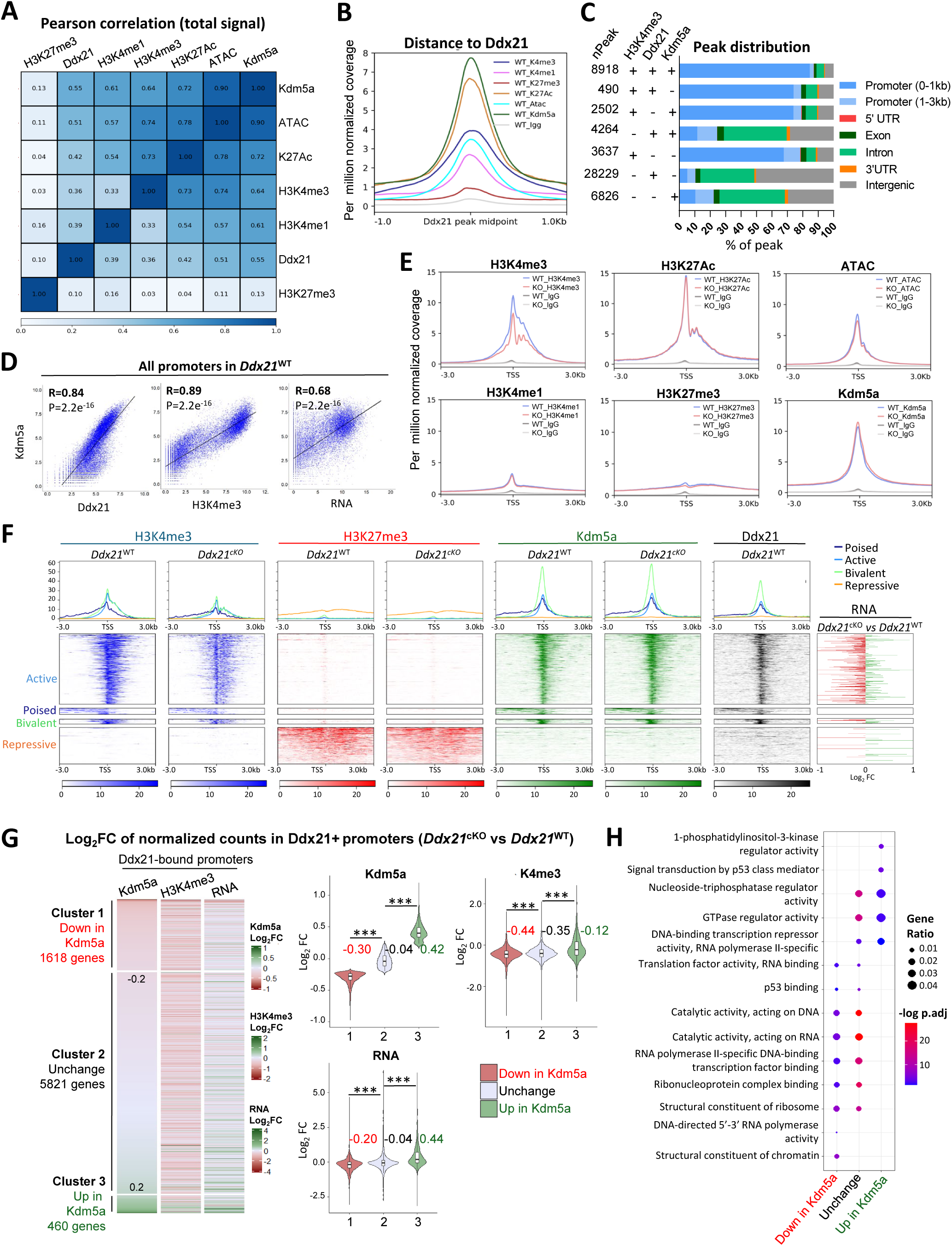
Ddx21 and Kdm5a co-occupy active promoters to preserve a transcriptionally active state.

To delineate the epigenetic alterations following Ddx21 knockout, we profiled the epigenetic landscapes in *Ddx21^cKO^* HSPC (Figures S11 and S12). A significant downregulation of H3K4me3 was observed at the transcription start site (TSS) following Ddx21 knockout (Figures 5E and S12). By contrast, H3K27ac and H3K4me1 were not markedly changed. Ddx21 loss had a mild effect on Kdm5a binding at TSS (Figure 5E). These changes suggest that Ddx21 may interact with Kdm5a to regulate gene transcription. Indeed, we found predominant occupancies for both Kdm5a and Ddx21 at TSS for active but not repressive genes; and Ddx21 knockout resulted in reduced expression of those active genes (as revealed by RNA expression) and reduced H3K4me3 peaks (Figure 5F). Interestingly, we found strong co-recruitment of Kdm5a/Ddx21 to active enhancers (Figure S13).

To understand genes regulated by Ddx21 and Kdm5a, we focused our analysis on the Ddx21-bound promoters. Gene clustering revealed 3 clusters based on Kdm5a binding changes upon *Ddx21* knockout (Figure 5G). In cluster-1 (1618 genes), Kdm5a binding was decreased (Log_2_FC < −0.2). Genes in this cluster were generally downregulated (low levels of RNA and H3K4me3) and were related to structural constituents of chromatin and ribosome (Figure 5H). In cluster 2, Kdm5a binding had marginal change but still have reduction in H3K4me3. In cluster 3 (460 genes), however, Kdm5a binding was increased (Log_2_FC > 0.2). Genes in this cluster were generally upregulated and were related to PI3K activity and p53 signal transduction. Together, these genomic analyses revealed the co-occupancy of Ddx21 and Kdm5a on active genes, consistent with their direct molecular interaction. Loss of Ddx21 altered the promoter-associated H3K4me3 level and gene expression.

### Epigenetic control of rDNA and cKit transcription by Ddx21 and Kdm5a

Given that DDX21 binds rDNA and coordinates rDNA transcription, we analyzed the genomic occupancy of Kdm5a at rDNA. In *Ddx21*^cKO^ HSPC, a marked decrease in H3K4me3 was observed at the spacer promoter (SpPr) region of rDNA, in line with the reduced rRNA transcription (Figures 6A and 3E). To test whether KDM5A modulates rDNA transcription, we treated K562 cells with KDM5A inhibitor CPI-455. As expected, CPI-455 treatment increased global H3K4me3 level, accompanied with elevated rRNA level and translational activity in DDX21-knockdown cells (Figure S15). Encouraged by these results, we tested whether CPI-455 could rescue the cellular defects in mouse HSPC. The result indicated that CPI-455 alone could enhance protein synthesis, but failed to rescue apoptosis and cell cycle arrest (Figures 6B, 6C and 6D). Interestingly, combined treatment with CPI-455 and PFT-α showed cooperative effect by further enhancing protein synthesis and meanwhile rescued apoptosis and cell cycle arrest in Ddx21-knockdown HSPC (Figures 6B, 6C and 6D). Next, we tested whether knockdowns of Kdm5a and/or p53 could rescue the defect in HSPC self-renewal by CFU assay. The result showed that Kdm5a knockdown could increase BFU-E number. Notably, dual knockdown of Kdm5a and p53 could significantly increase BFU-E but slightly on CFU-GM numbers (Figures 6E and S16). Lastly, we investigated the effect of dual inhibition of Kdm5a and p53 on fetal HPSCs *in vivo*. Injections of CPI-455 and PFT-α to pregnant mice at E11.5-E12.5 resulted in increased expression of *cKit*, as analyzed for the E13.5 FL cells by flow cytometry. As a result, both LS^−^K and LSK HSPC populations were increased (Figure 6F). These data suggest a partial rescue of HSPC survival/proliferation and translational capacity by suppressing both Kdm5a and p53.

**Figure 6.**
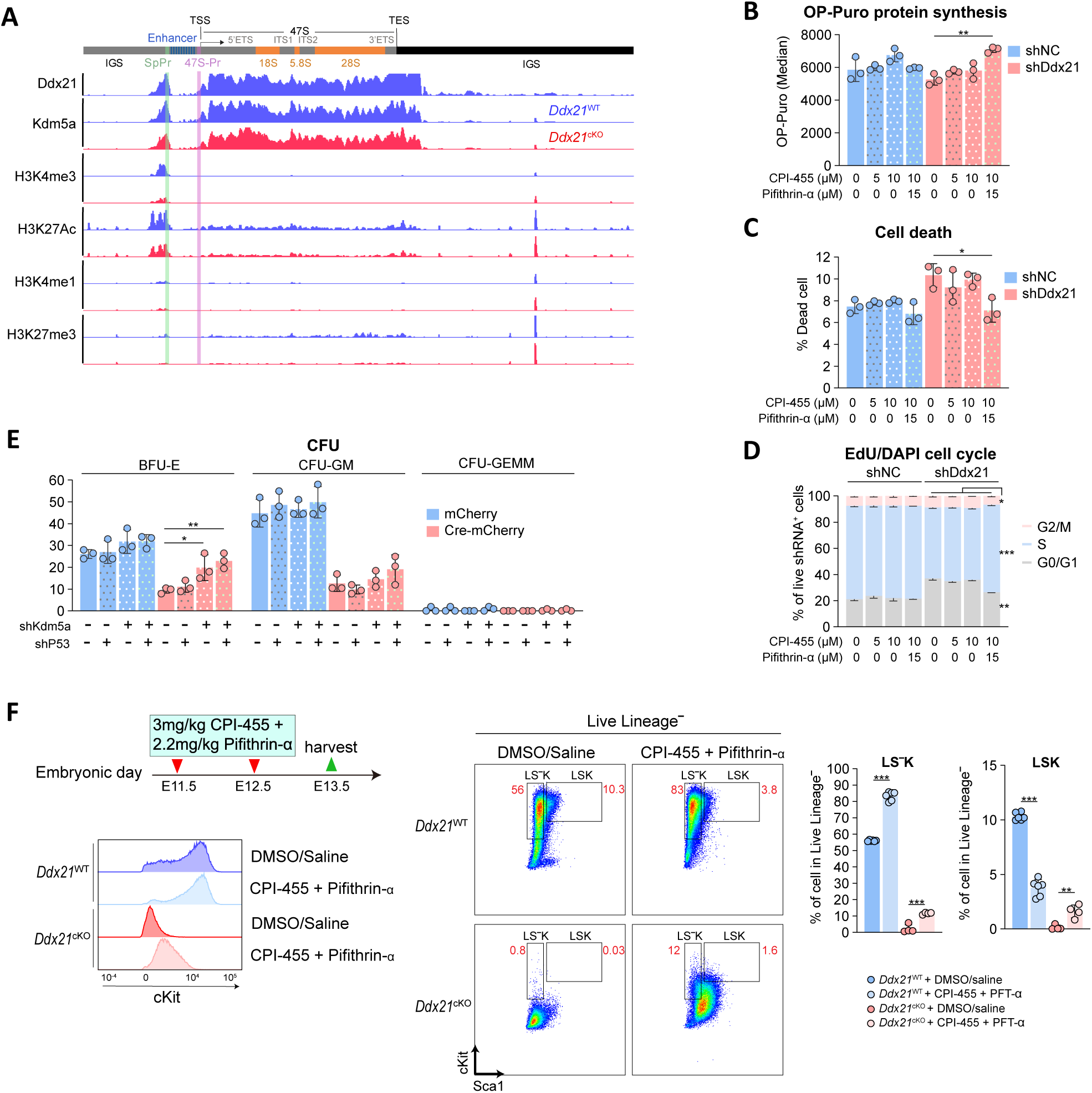
Dual inhibition of Kdm5a and p53 restores rRNA level and translational capacity in Ddx21-deficient hematopoietic progenitors.

To summarize, our work supports a working model where DDX21 is an indispensable ribosome biogenesis factor for supporting fetal HSPC survival and expansion. Mechanistically, DDX21 interacts with KDM5A to epigenetically maintain an active chromatin state that enables active transcription of hematopoietic genes (e.g. *cKit* and *Gata1*) and rDNA, the perturbation of which triggers loss of ribosome homeostasis, p53-induced cell death and cell cycle arrest, leading to hematopoietic failure.

## DISCUSSION

This study unveils an epigenetic role of RNA helicase Ddx21 in safeguarding fetal hematopoiesis by collaborating with histone modification enzyme. We demonstrate that Ddx21 loss results in downregulation of H3K4me3 associated with active promoters. As a result, rDNA transcription and ribosome biogenesis decline, activating p53-dependent stress responses, and impair hematopoietic progenitor proliferation and self-renewal, ultimately leading to HSPC depletion and fetal anemia. Importantly, we identify a mechanism whereby Ddx21 partners with Kdm5a to maintain an active chromatin state at ribosomal and hematopoietic genes, ensuring robust transcriptional output necessary for HSPC expansion. These findings bridge ribosome biology with epigenetic regulation, offering new insights into the pathogenesis of ribosomopathies.

The link between defective ribosome biogenesis and hematopoietic failure is well-established in ribosomopathies such as DBA, which manifests fetal or neonatal anemia due to low erythroid precursors (Da Costa et al., 2020; Iskander et al., 2021; Ulirsch et al., 2018). However, most studies focus on ribosomal protein mutations, leaving non-structural ribosomal factors underexplored. Our work positions Ddx21 as a key non-structural regulator whose loss phenocopies ribosomopathy-associated anemia, underscoring the vulnerability of HSPCs to perturbations in ribosome supply. The observed p53 activation aligns with the canonical “ribosomal stress checkpoint” model, where ribosome insufficiency triggers p53-mediated cell cycle arrest (Hubackova et al., 2020; Iskander et al., 2021; Zhang et al., 2003).

The discovery of Ddx21-Kdm5a interaction expands our understanding of how ribosome biogenesis intersects with chromatin dynamics. Kdm5a, a H3K4me3 demethylase, is typically proposed to repress transcription by removing active histone marks (Beshiri et al., 2012; Kong et al., 2018; Xu et al., 2021). Intriguingly, other studies indicate a transcripitonal activator role of Kdm5a in controlling cell cyle progression when interacting with Rb (Benevolenskaya et al., 2005; Brier et al., 2017). The specific role of Kdm5a in the epigenetic control of ribosome biogenesis remains unknown, although other members of the Jumonji family (e.g. KDM2A/B) have been shown to repress rDNA transcription (Frescas et al., 2007; Tanaka et al., 2010). Our data suggest that Ddx21 may sequester Kdm5a at active promoters, limiting its demethylase activity and preserving a high H3K4me3 level. This model is supported by the co-occupancy of Ddx21 and Kdm5a at H3K4me3-rich promoters and the restoration of rRNA expression upon Kdm5a inhibition. This mechanism may explain why rRNA, which require high transcriptional activity, are particularly sensitive to Ddx21 loss. Notably, this interaction appears RNA-dependent, as RNase treatment impaired Ddx21-Kdm5a interaction, suggesting that RNA scaffolds or nascent transcripts stabilize their association at active loci. Profiling of Ddx21-associated chromatin RNA may help to address this question.

Therapeutic implications emerge from the partial rescue of HSPC defects by targeting Kdm5a and p53 using preclinical inhibitors. While p53 blockade alleviates apoptosis and, as shown in zebrafish, could rescue morphological phenotypes associated with ribosomopathy but not erythroid defects (Boglev et al., 2013; Yadav et al., 2014), restoring ribosome biogenesis via epigenetic modulation may be necessary to better rescue hematopoiesis. This aligns with clinical observations in DBA, where glucocorticoids improve anemia but fail to address ribosomal defects (Maceckova et al., 2022; Vlachos and Muir, 2010). Targeting the DDX21-KDM5A complex could offer a complementary strategy, though further studies are needed to optimize such approaches.

In conclusion, our findings redefine Ddx21 as a multifunctional hub integrating ribosome biogenesis with epigenetic regulation. By coupling rRNA synthesis to chromatin modification, Ddx21 ensures that fetal HSPCs meet the extraordinary biosynthetic demands of the developing fetus, especially for the predominant erythroid lineage. Disruption of this axis not only illustrates the etiology of ribosomopathies but also opens avenues for therapies targeting the epigenetic-ribosome interface.

### Limitations of study

Limitations of this study include the reliance on murine models and cultured HSPC systems, which may not fully recapitulate human pathophysiology. Additionally, the precise molecular cues directing DDX21-KDM5A complex to specific genomic loci remain unclear. Future work should explore whether other chromatin modifiers or RNA-binding proteins collaborate with Ddx21 to fine-tune transcriptional programs in HSPCs. Thirdly, ablation of Ddx21 in adult HSPC and its effect on adult hematopoiesis remain unknown. Furthermore, whether heterozygous knockout (*Vav1-Cre;Ddx21^f/+^*) would develop blood disorder (e.g. myelodysplasia) in aged mice has not been examined, altough current data indicated that these mice were healthy and comparable with WT. Last but not the least, due to the cellular heterogenicity of HSPC population, the transcriptional readout of LK cells was biased towards the erythroid progenitors (MEP). Simutaneously profiling of the transcriptome and epigenetome of HSPC at single cell level would provide a higher resolution of the hematopoietic atlas.

## METHODS

### Animals and embryo analysis

*Ddx21^f/f^* mice were custom-generated by Cyagen with insertion of loxP sites into introns flanking exon 3-5, creating a nonsense mutation upon Cre-mediated recombination. B6.Cg-Commd10Tg(Vav1-icre)A2Kio/J (*Vav1-Cre*) (JAX #008610), B6.SJL-Ptprca Pepcb/BoyJ (CD45.1) (JAX #002014) and Gt(ROSA)26Sortm4(ACTB-tdTomato,-EGFP)Luo/J (*mTmG^f/f^*) (JAX #007676) mice were imported from the Jackson Laboratory. All mice were maintained in C57Bl/6 background. Mice were bred and housed in pathogen-free animal facilities with regular light-dark cycle. Embryo was determined as day0.5 when vaginal plug is observed on the following day after mating. Both male and female were studied and randomly assigned for experiments, except transplantation recipients which are male-only. Genotyping and Ddx21-KO efficiency PCR was performed with RapidTaq (Vazyme), primer sequences are listed in Supplemental Table 1. All animal experimental protocols were reviewed and approved by Animal Experimentation Ethics Committee of CUHK.

### Fetal liver HSPC cell culture

Live LK cells from fetal liver were FACS-sorted and cultured in StemPro™-34 SFM (Gibco) with SCF (100ng/ml), FLT3L (50ng/ml), TPO (50ng/ml), Il3 (10ng/ml), Il6 (10ng/ml), L-glutamine (1x) and Anti-Anti (1x), maintained at density of < 0.2M cells/ml. Lentiviral transduction was performed on day 1, where each type of lentivirus was added at MOI=10 in 8µg/ml polybrene. Cells were centrifuged at 1000g 37 1hr and incubated for 5hrs, full medium change was performed afterwards with addition of drug/DMSO as indicated. Respective control lentivirus (shNC-GFP/mCherry or mCherry-only lentivirus) was added to certain samples to ensure each sample was subjected to equivalent total lentivirus MOI. After 3 days of transduction, live shRNA^+^ cells were analyzed or FACS-sorted.

### Flow cytometry and cell sorting

All antibody staining was carried out in 1%FBS/PBS for 30min 4. Centrifugation was performed in 500g 5min 4 or 800g after permeabilization. Antibodies are listed in supplemental table 3 and table 4. Flow cytometry was performed on either BD Symphony A5 SE or BD FACSDiscovery S8. Analysis was performed using FlowJo v10.8.1. Cell sorting was performed with either BD FACSdiscovery S8 or BD FACSAria Fusion in purity mode. Sample and collection tube were maintained at 4°C. Gating strategies were illustrated in Figures S2 and S4A-B.

For fetal liver HSPC analysis, HSC isolation and analysis of transplanted bone marrow, cells were stained with lineage-biotin Ab (CD3, B220, CD11b, Gr1, Ter119) and viability dye. After PBS wash and centrifuge, cells were stained with streptavidin, cKit, Sca1, CD48, CD150, CD16/32 and CD34. CD45.1/CD45.2 was also included for transplantation with changes in fluorophore conjugates. Flow cytometry was performed after wash and resuspension. For erythroid fraction, Ter119 was omitted in lineage-biotin panel, cells were subsequently stained with CD71 and Ter119. For leukocytes, cells were stained with viability dye, CD11b, Gr1, CD3 and B220. For sorting of fetal liver LK cells, cells were stained with cKit after lineage-biotin panel staining. For peripheral blood analysis of transplanted mouse, leukocytes were stained with CD3, B220, CD11b and viability dye with CD45.1/CD45.2 or CD45 depending on the transplantation.

Human HSPC was analyzed or sorted after staining with CD34, CD43, CD45 and viability dye. For apoptosis, Apotracker™ Green (Biolegend) was co-stained with surface markers. EdU and OP-puromycin detection by Click-iT™ (Life Technologies) was performed after surface staining, fixation and permeabilization.

### Colony formation assay (CFU)

Mouse hematopoietic colony formation assay was performed using MethoCult™ GF (M3434, Stemcell), colonies were counted on day12. For fresh fetal liver cells, 20000 cells were seeded in triplicates. For lentiviral transduced fetal liver cells, FACS-sorted LK cells were cultured and transduced as mentioned above, 6000 live transduced cells were FACS-sorted and seeded in duplicates. Human hematopoietic colony formation assay was performed using MethoCult™ GF (H4536, Stemcell), colonies were counted on day16. 3000 lentiviral transduced HSPC as described above were seeded in duplicates.

### Fetal liver and bone marrow cell isolation

Isolated tissues and cells were kept at either on ice or at 4 throughout the entire process of harvesting, centrifugation and cell sorting. Dissected fetal livers were washed twice with PBS and kept in fetal liver juice (2% FBS/1mM EDTA/10mM Glucose/PBS) on ice. Fetal livers were triturated about 20 times with P1000 pipette. Cell suspension was strained with a sterile 70µm cell strainer (Beyotime) and kept in fetal liver juice for no longer than 1 hour until subsequent experiments.

For bone marrow isolation, tibia and femur were harvested. Bones were stripped clean of connective tissues and muscles; epiphysis were removed at both ends. Bone marrow was flushed with staining buffer (1% FBS/PBS) with 25G needle and strained with a sterile 70µm cell strainer (Beyotime). After centrifugation, cells were resuspended in 100µL RBC lysis buffer (150mM NH_4_Cl/10mM NaHCO_3_/1mM EDTA) per bone for 2min on ice and quenched with staining buffer. After centrifugation, cells were strained again with 70µm strainer.

### Transplantation

All recipient mice were C57Bl/6J male of age 2-3 months old in either WT or CD45.1 background and kept in IVC cages. They were fed with 0.5mg/ml enrofloxacin from 3 days before to 14 days after transplantation. For complete myeloablation, recipients were subjected to 10Gy total body irradiation once during the day of transplantation using Gammacell 3000 irradiator with ^137^Cs radiation source. For each recipient, 200uL cell suspension in HBSS were transplanted via tail vein injection using a 30G needle. The time between irradiation and injection was kept within 3-6 hours.

### Cell lines

All cells were maintained in 37, 5% CO_2_. 293T cells were cultured in high glucose DMEM (Life Technologies) supplemented with 10% FBS, 1x NEAA, 2mM glutamine, 1mM Sodium pyruvate and 1x Antibiotic-Antimycotic. K562 cells were cultured in IMDM (Life Technologies) supplemented with 10% FBS, 1x NEAA and 1x Antibiotic-Antimycotic. H1 hESC were cultured in mTESR+ medium on Geltrex coated wells, medium was changed every 1-2 days. H1 was maintained at < 60% confluency and passaged using Dispase and scraping followed by 2-3 times of trituration. Expi293F™ Cells were maintained in cells were cultured in SMM 293-TIL-N complete medium (SinoBiological) and shaken in 250ml flasks at 150rpm, maintained at <5×10^6^ cells/ml. 293T and K562 were frozen in 90%medium/10%DMSO, H1 cells in StemCell Keep (Funakoshi). K562 cell lentiviral transduction was performed at MOI=5 with 8µg/ml polybrene and incubated overnight. Full medium change was performed on the following day.

### Differentiation of CD34+ HSPC from hESC

hESC H1 (WiCell) was differentiated into HSPC using STEMdiff™ Hematopoietic Kit (#5310, Stemcell). On day10, 1×10^6^ functional titre of shRNA-mCherry lentivirus was added to each 12-well with 8ug/ml polybrene and incubated for 5 hours. Full medium change was performed afterwards with addition of drug/DMSO as required. After 2 days, HSPC (live CD45^+^ CD34^+^ CD43^+^ shRNA-mCherry^+^) were FACS-sorted or analyzed.

### Lentiviral packaging, purification and titre determination

Second generation lentivirus was generated as previously described with some modifications (Tiscornia et al., 2006). In brief, 22×10^6^ 293T cells in 15-cm dish were transfection with total 55µg of plasmids [packaging plasmids pMD2.G (Addgene #12259), psPAX2 (Addgene #12260) and transfer plasmid] in 1:1:1.5 molar ratio with 140µg PEI (1:2.5). Medium was replaced on day1, and viral medium collected on day2 and day3. Cell debris was removed by performing 1000g 5min 4 centrifugation twice, followed by filtering the supernatant with 0.45µm PES filter. Virus was concentrated with 20000g 2hr 4 centrifugation. Viral pellet was resuspended in DMEM, aliquoted and kept at −80. Virus was freeze-thawed for no more than 3 times. Cre-IRES-mCherry lentivirus was purchased from BrainVTA. Functional lentiviral titre was calculated by serial transduction of 293T cells in 8ug/ml polybrene and calculating the percentage of fluorescence positive cells after 3 days of transduction. shRNA sequences were listed in Supplemental Table 2.

### EdU/DAPI cell cycle flow cytometry

5-Ethynyl-2’-deoxyuridine (EdU) and 4’,6-Diamidino-2-Phenylindole, Dihydrochloride (DAPI) cell cycle analysis was performed using Click-iT™ Plus EdU Alexa Fluor™ 647 Flow Cytometry Assay Kit (Life Technologies) according to product manual. Cells were incubated in 10µM EdU at culture condition for 1hr. Pregnant mice were injected i.p. with 30ug/g EdU, fetal liver was harvested 1.5hr post injection. After surface panel staining, fixation, permeabilization and Click-iT reaction, cells were stained in 1ug/ml DAPI for 30min at 4. Events were captured in linear mode with flow rate <1000 events/sec for all samples.

### O-propargyl-puromycin (OP-Puro) nascent protein synthesis flow cytometry

Protein nascent protein synthesis was determined using Click-iT™ Plus OPP Alexa Fluor™ 647 Protein Synthesis Assay Kit (Life Technologies) according to product manual. Cells were incubated in 10µM OP-Puro for 30min in culture conditions. Click-iT reaction was performed after surface panel staining, fixation and permeabilization. 1X Click-iT saponin-based permeabilization and wash reagent from Click-iT™ Plus EdU Alexa Fluor™ 647 Flow Cytometry Assay Kit was used instead of the suggested Triton-X based permeabilization buffer.

### 5-ethynyluridine (EU) nascent RNA transcription

Cells were incubated in 10µM 5-ethynyluridine (EU) for 1hr in culture conditions. RNA was extracted by Total RNA was extracted by FastPure Cell/Tissue Total RNA Isolation Kit V2 (Vazyme). Click-iT reaction was performed with 3µg RNA and 0.25mM Biotin-PEG4-Azide (Beyotime) in 300µL total volume for 30min RT using BeyoClick kit (Beyotime). Total RNA after click-iT reaction was extracted again by the same extraction kit in 30µL eluate. 50µL streptavidin bead (Beyotime) was washed with 1xTBS twice, followed by 0.05M NaCl (DEPC-treated) twice and 0.1M NaCl (DEPC-treated) once. Beads were resuspended in 1x bind and wash buffer (5mM Tris-HCl pH7.5, 0.5mM EDTA, 1M NaCl, 0.01% Tween-20) and mixed with the RNA eluate for 1hr RT with end-to-end rotation. Beads were washed twice in 1x bind and wash buffer and resuspended in 40µL PrimeScript™ RT Master Mix (Takara). Reverse transcription was carried out on bead in 37 for 1hr followed by 85 5min to inactivate the enzyme and release the cDNA. For qPCR, cDNA was diluted 10-fold and 4µL was used per 10µL qPCR reaction using TB Green® Premix Ex Taq™ II (Takara).

### Reactive oxygen species (ROS) detection by flow cytometry

Flow cytometry ROS quantification by ROS dye H_2_DCFDA (Life Technologies) was performed as previously described (Bode et al., 2020). In brief, cells were incubated in 50µL 10µM H_2_DCFDA/PBS 37 20min after surface panel staining and washing. Cells were then diluted with 250 µL staining buffer and kept on ice. They were subjected to flow cytometry immediately. Signal was detected in with blue laser in filter BP537/32 “GFP/FITC channel” of BD FACS Symphony A5 SE.

### Peripheral blood leukocyte preparation and staining

Tail vein peripheral blood was extracted by tail nicking with 27G needle. 20µL blood was mixed with 0.5µL 0.5M EDTA, pH8.0 and incubated with 0.5µL TruStain FcX (Biolegend) for 10min. RBC was lysed by adding 200µL RBC lysis buffer (150mM NH_4_Cl/10mM NaHCO_3_/1mM EDTA) for 10min and quenched with 1ml PBS. Leukocytes were pelleted with 200g 5min centrifuge, stained with surface Ab panel and assessed by flow cytometry for chimerism analysis.

### Hemoglobin quantification assay

Hemoglobin content in fetal liver was measured with Hemoglobin Assay Kit (Colorimetric) (Abcam ab234046) in duplicates according to product manual. Each reaction consisted of 0.24 x 10^6^ total fetal liver cells or hemoglobin standard (Sigma H2500) resuspended in 20µL ddH_2_O and 180µL detection buffer. Readout measured at 575nm using a plate reader.

### CPI-455 and Pifithrin-α injection

Stock CPI-455 (25mM, 6.96µg/µL) was prepared by dissolving in DMSO. Stock pifithrin-α (PFT-α) (10mM, 3.67µg/µL) was prepared in DMSO. Both CPI-455 and PFT-α were freshly diluted in saline solution (0.9%NaCl) to final 100uL injection volume. CPI-455 and PFT-α dosage was 3mg/kg and 2.2mg/kg respectively. Pregnant mice were injected intraperitoneally at E11.5 and E12.5, fetal liver harvested at E13.5. Control mice were injected with same equal injection volume and ratio of saline/DMSO mix.

### Raising of polyclonal antibody against mouse Ddx21

Antibody targeting mouse Ddx21 was raised against M719-Q851 peptide of mouse Ddx21 (Q9JIK5). The peptide has low sequence similarity with other proteins including other Ddx family proteins. Rabbit Ddx21 antibody was produced and purified by ABclonal. Specificity was tested by immunofluorescence staining in *Ddx21*^cKO^ fetal liver LK cells.

### Fetal liver cell cytospin, RNase A treatment and immunofluorescence

RNase A treatment of cells for immunofluorescence was performed as previously described with some modifications (Beltran et al., 2016). In brief, 20000 fetal liver LK cells were permeabilized with 0.05% Tween-20/PBS for 10min on ice, washed with 1ml PBS, centrifuged and resuspended with 10 µL of 1mg/ml RNase A/PBS for 30min RT. Cell suspension was diluted to 100 µL with PBS and centrifuged at 40g 3min in a cytospin centrifuge to prepare the cytospin slides. Cells were immediately fixed with 4% PFA/PBS for 10min, blocked and permeabilized in 0.5%Triton-X/10% goat serum/PBS for 30min and incubated in primary ab in 0.1%Tween-20/5% Goat serum/PBS at 4°C O/N. After 3 washes in PBS, respective secondary antibodies (1:1000) and 1ug/ml Hoechst 33258 (Life Technologies) in 0.1%Tween-20/5% Goat serum/PBS were applied for 1hr RT. Slides were mounted with ProLong™ Glass Antifade Mountant (Life Technologies). Antibodies used were listed in Supplemental Tables 3-5.

### PLA proximity ligation assay staining

PLA staining on the fetal liver LK cell cytospin slides was performed using Duolink® In Situ Orange Starter Kit Mouse/Rabbit (Sigma) according to product manual. In brief, cytospin slides were fixed in 4% PFA 10min RT and permeabilization in 0.1%Triton-X/PBS for 10min RT. Blocking and subsequent procedures was performed following the product manual. Negative control was performed by omitting one of the Ab (only Ddx21 Ab was added). Antibodies used were listed in Supplemental Tables 3-5.

### Imaging and quantification

Immunofluorescence and proximity ligation assay staining were imaged and visualized with either of the following microscope/software pair: Nikon Ti2-E widefield fluorescence microscope with NIS-Element Viewer, Olympus FV1200 confocal microscope with FV10-ASW or Leica SP8 confocal microscope with LAS X. The same imaging configurations and LUTs were applied to all images in the same batch of staining. Representative images were slightly enhanced with “brightness/contrast” layer in Photoshop.

For signal quantification, nuclear masks were made using DAPI signal by ImageJ (Threshold -> Watershed -> ROI). Only nucleus with proper nuclear morphology (no smear or fold) were quantified to neglect damaged nucleus due to cytospin. For every nucleus in the mask, total relative fluorescence unit (RFU) was measured as: [Area of nucleus x (Nuclear mean grey value - background mean grey value)] using ImageJ (Measure: area, mean grey value). Total RFU of different targets were multiplied to different constants for better visualization of the quantification.

### Plasmids

Lentiviral packaging plasmids pMD2.G (#12259), psPAX2 (#12260), Flag-DDX21 overexpression plasmid (#128803), SIN3A overexpression lentiviral plasmid (#142686), HDAC1-V5 overexpression lentiviral plasmid (#144813), HA-KDM5A overexpression plasmid (#14799), eGFP-TwinStrepTag plasmid (#141395) and BioID2-HA construct (#74224) were purchased from Addgene. TwinStrepTag from #141395 was subcloned to N-terminal of SIN3A in the overexpression lentiviral plasmid (#142686), creating TwinStrepTag-SIN3A overexpression lentiviral plasmid. Full length HDAC2-V5 was cloned from H1 hESC RNA using 5’-tttaatcgatATGGCGTACAGTCAAGGAGGC3’ and 5’-tttaacgcgtctacgtagaatcgagaccgaggagagggttagggataggcttaccGGGGTTGCTGAGCTGTTCTG-3’, replacing HDAC1-V5 in HDAC1-V5 plasmid, creating HDAC2-V5 overexpression lentiviral plasmid.

BioID2-HA construct from #74224 was subcloned to pLVX backbone, then full length DDX21 sequence from #128803 or SV40NLS sequence was subcloned to N-terminus of BioID2-HA to create fusion BioID2 lentiviral plasmids. Lentiviral shRNA transfer plasmid were obtained by cloning shRNA sequences to pLVX-U6-mCherry/GFP plasmids (Homemade). shRNA sequences were listed in supplemental table 2.

### Purification of DDX21 protein

The full-length human DDX21 gene was cloned into a pET-N-6xHis-TEV (Tobacco etch virus) vector. The construct was then transformed into BL21-CodonPlus (DE3)-RIL competent cells for expression. After expression, the cells were harvested by centrifugation and resuspended in a buffer containing Tris (pH 8.0) and 500 mM NaCl. Cell lysis was performed using a high-pressure homogenizer, followed by affinity purification using a HisTrap HP column (Cytiva). The purified DDX21 protein was concentrated to a final concentration of 10 mg/mL and stored at −80°C for subsequent experiments.

### SDS-PAGE Western blot

Cell lysates were prepared by digestion in Lysis Buffer (Beyotime, P0013C) for 10min on ice. Supernatant was extracted after 16000g 10min 4 centrifugation. Protein concentration was measured by Pierce™ BCA Protein Assay Kits (Life Technologies). Lysate was mixed with 4x Laemmli Sample Buffer (Bio-rad) and incubated at 95 for 10min. Proteins were resolved in 8-15% SDS-PAGE gels (Vazyme) depending on target protein size. After electrophoresis, gels were transferred to 0.22µm PVDF membrane by either overnight 4 methanol based wet transfer method or by semi-dry transfer using Trans-Blot® Turbo™ Transfer System (Bio-rad). Membrane was blocked in 5%BSA/PBS for 1hr and incubated in primary ab overnight 4. After 4 washes in 1%Tween/PBS, secondary ab was applied and incubated for 1hr RT. After 4 washes in 1%Tween/PBS, membrane was covered in Pierce™ ECL Western Blotting Substrate (Life Technologies) and visualized with ChemiDoc MP Imaging System (Bio-rad).

### Co-Immunoprecipitation

Cell lysates were extracted with the same method in western blot. Immunoprecipitation was performed with or without RNase A treatment. A final concentration of 300μg/mL RNase A (Sigma, 10109142001) was used to treat cell lysates. Cell lysates were incubated with respective beads at 4 °C for 12 h. The beads were subsequently washed twice with the wash buffer (Life Technologies, 87787) and once with UltraPure water (Life Technologies). Beads with binding components were boiled in 1× SDS-PAGE sample loading buffer (Beyotime, P0015A) for 10min, and the eluted proteins were resolved by SDS-PAGE Western blot. Antibodies and beads were listed in Supplemental Tables 3-5.

### Recombinant protein pull-down assay

KDM5A or HDAC1 were incubated with equimolar DDX21 protein, anti-DDX21 antibody and Dynabeads Protein G in 500μl of binding buffer containing 25mM Tris-HCl, 150mM NaCl, 1mM EDTA, 1% NP-40 and 5% glycerol, pH7.4, at 4 for 12hr. The beads were washed and boiled, the complex was ressolved by SDS-PAGE western blot. Recombinant DDX21 protein was purified as above, KDM5A (BPS Bioscience, 50110) and HDAC1 (BPS Bioscience, 50051) was purchased.

### RNA extraction and quantitative reverse transcription PCR (qRT-PCR)

Total RNA was extracted by FastPure Cell/Tissue Total RNA Isolation Kit V2 (Vazyme) or Direct-zol RNA Microprep kit (Zymo Research) with gDNA removal. Reverse transcription and qPCR were performed using PrimeScript™ RT Master Mix (Takara) and TB Green® Premix Ex Taq™ II (Takara) respectively. cDNA was diluted 20-fold and 2µL was used for each 10µL qPCR reaction. Each target was performed in triplicates on QuantStudio 7 Flex (Applied Biosystem) in “fast” mode, other settings in default. Cycle threshold values were converted to fold change expression by 2^-ddCT^ using Gapdh as control. qPCR primer sequences were listed in Supplemental Table 6.

### TapeStation rRNA profiling

rRNA quantification and profiling was performed using RNA ScreenTape Analysis or High Sensitivity RNA ScreenTape (Agilent). 18S, 28S rRNA and total RNA concentrations were determined accordingly after automatic peak assignment. For samples to be compared with, equal number of cells were used for RNA extraction and the extraction was performed at the same time using the same method and kit.

### DDX21-BioID2 proximity protein pulldown and mass spectrometry (MS)

BioID2 detection of protein interactome was performed as previously described with some modifications. K562 cells were transduced with lentivirus carrying DDX21-BioID2-HA or SV40NLS-BioID2-HA. Nuclear localization of the fusion BioID2 proteins were verified by HA immunofluorescence staining. After 5 days of transduction, cells were incubated in 50µM D-biotin (Sigma) for 16 hours in culture condition in triplicates. 30M cells were lysed in 1ml fresh lysis buffer [50mM NaF, 100mM NaCl, 5mM EDTA, 50mM HEPES, 0.2%SDS, 0.5% IGEPAL, 1mM PMSF, 1.5mM Na_3_VO_4_, 1mM DTT, 1x protease inhibitor cocktail (Roche) and 1µL of 30U/µL benzonase (Sigma E1014) for 10min on ice. Lysate was pulse sonicated at 50% Amplitude (1sec On, 1sec Off) for 5min using Branson SLPe (2.4mm tip diameter). After 16000g 10min 4 centrifugation, the supernatant was diluted to 0.1% SDS. Protein concentration was determined by BCA, 15mg protein equivalent of lysates were incubated with 1ml streptavidin beads (Beyotime) O/N at 4 with end-to-end rotation. Beads were washed with wash buffer 1 twice and wash buffer 2-4 once for 5min each with end-to-end rotation [Wash buffer1: 2% SDS; wash. Wash buffer 2: 0.1% sodium deoxycholate, 1% Triton X-100, 500 mM NaCl, 1 mM EDTA, 50 mM HEPES pH 7.5. Wash buffer 3: 10 mM Tris-HCl pH 8.0, 250 mM LiCl, 1 mM EDTA, 0.5% IGEPAL, 0.5% sodium deoxycholate. Wash buffer 4: 50mM Tris-HCl, pH8.0].

Bead-protein were frozen and sent for 4D-Data Independent Acquisition (DIA) label-free quantitative LC-MS/MS analysis (Omicsolution). Beads were washed in 1ml 50mM NH4HCO3. After centrifugation, beads was reduced with 2µL of 0.5M Tris 2-carboxylethyl phosphine (TCEP) 37 60min and alkylated with 4µL 1M iodoacetamide (IAM) for 40min RT. Five fold volume of cold acetone were added to precipitate the proteins at −20 overnight. After centrifugation at 12000g 20min 4°C, the pellet was washed twice by 1mL of pre-chilled 90% acetone aqueous solution and re-suspended with 100µL of 100mM triethylammonium bicarbonate (TEAB) buffer. Trypsin was added at 1:50 trypsin-to-protein mass ratio and incubated at 37 overnight, followed by desalting with Pierce C18 Spin Tips, and then dried in a speed vacuum concentrator.

The UltiMate 3000 (Thermo Fisher Scientific, MA, USA) liquid chromatography system was connected to the timsTOF Pro 2, an ion-mobility spectrometry quadrupole time of flight mass spectrometer (Bruker Daltonik). Samples were reconstituted in 0.1% FA and 200 ng peptide was separated by AUR3-15075C18 column (15 cm length, 75 µm i.d, 1.7 µm particle size, 120 Å pore size, IonOpticks) with a 30 min gradient starting at 2.2% buffer B (80% ACN with 0.1% FA) followed by a stepwise increase to 28% in 16min, 38% in 8min, 90% in 3min and stayed there for 3min. The column flow rate was maintained at 400nL/min with the column temperature of 50°C. DIA data was acquired in the diaPASEF mode. We defined 22 × 40 Th precursor isolation windows from m/z 349 to 1229. To adapt the MS1 cycle time, we set the repetitions to variable steps (2-5) in the 13-scan diaPASEF schemein our experiment. During PASEF MSMS scanning, the collision energy was ramped linearly as a function of the mobility from 59 eV at 1/K0 = 1.6 Vs/cm^2^ to 20 eV at 1/K0 = 0.6 Vs/cm^2^.

Proteomic spectral analysis and quantification was performed by MS service provider using Spectronaut 19 (Biognosys) with default settings. Database used was Homo Sapiens (Ver. 2024, 20608 entries) from uniport. Trypsin was the digestion enzyme and specific was the digest type. Carbamidomethyl on cysteine was specified as the fixed modification. Oxidation on methionine, Acetyl on protein N-term were specified as the variable modifications. Retention time prediction type was set to dynamic iRT. Data extraction was determined by Spectronaut based on the extensive mass calibration. Spectronaut will determine the ideal extraction window dynamically depending on iRT calibration and gradient stability. Qvalue (FDR) cutoff on precursor, peptide and protein level was 1%. Decoy generation was set to mutated which is similar to scrambled but will only apply a random number of AA position swamps (min=2, max=length/2). Normalization strategy was set to Local normalization. Peptides which passed the 1% Q value cutoff were used to calculate the major group quantities with MaxLFQ method.

To obtain DDX21-specific affinity protein list, Spectronaut output was further analysed using MSstats. By comparing proteins obtained from DDX21-BioID2 to that of NLS-BioID2, significant proteins (q<0.05) were considered as DDX21 affinity proteins.

### RNA-seq and data analysis

Biological triplicates (embryos/culture wells) were collected for each transcriptomic dataset. Total RNA was extracted by Direct-zol RNA Microprep kit (Zymo Research) with gDNA removal. Enrichment of mRNA, construction of cDNA library (ABclonal RK20350) and PE150 sequencing (NovaSeq 6000) was performed by HaploX. Reads were aligned to hg38 or mm10 using STAR aligner (2.7.11b) [--outFilterMultimapNmax 20--alignSJoverhangMin 8 -- alignSJDBoverhangMin 1--outFilterMismatchNoverReadLmax 0.04--alignIntronMin 20 -- alignIntronMax 1000000 -- alignMatesGapMax 1000000 --outFilterType BySJout --sjdbScore 1]. Count matrix was constructed by paired read counting of exons using subread:featureCounts (2.0.8). To remove low expressing genes, genes where less than half of the samples have EdgeR:CPM >1 were filtered out. Normalized counts were obtained by DEseq2:RLE (1.44.0). Gene name/ID conversion was performed using AnnotationDbi (1.66.0). Genes with Log2FC > 1 and p.adj < 0.05 were used for subsequent differential analysis. GO/KEGG enrichment and GSEA was performed using clusterProfiler (4.12.6) and pathview (1.40.0). GO term clustering and visualization was performed using Cytoscape (3.10.3). Volcano plot and heatmap were generated using EnhancedVolcano (1.22.0) and ComplexHeatmap (2.20.0) respectively. Raw and processed data files were deposited in the Gene Expression Omnibus (GEO) under accession number GSE296783.

### CUT&Tag and data analysis

CUT&Tag libraries were constructed in biological duplicates using CUTANA™ CUT&Tag Kit (Epicypher) according to product manual. 30000-60000 FACS-sorted live fetal liver LK cells were used per library with 15-18 cycle of PCR amplification. Libraries were validated by Agilent high sensitivity dna kit, PE150 sequencing (NovaSeq X) was performed by HaploX. QC was performed with FastQC (0.12.1). Adapter and 3’ polyG was removed with fastp (0.23.4) -g -- detect_adapter_for_pe. Reads were aligned to UCSC mm10 genome using bowtie2 (2.5.4) (--local --very-sensitive --no-mixed --no-discordant --phred33 --threads 8 -I 20 -X 1000). Alignments with MAPQ <30 (including unmapped and multi-mapped regions) and unwanted regions/chromosomes (chrM, random, chrUn, chrEBV) were filtered out using samtools (1.20.0). For H3K27Ac, duplicate reads were removed with picard (3.2.0). BAM files were converted to BEDPE using bedtools (2.31.1). Broadpeaks were called using macs2 (2.2.9.1) with FDR cutoff of q<0.1 to 0.01 depending on S/N ratio. Peaks from replicates were merged whenever there is 1bp overlap by bedtools. Normalized and scaled to per million bigWig tracks were generated using deepTools:bamCoverage (3.5.5) with scaling factor calculated as 1000000/(EdgeR:TMM scale factor on reads in 10000bp background bin * library size). Blacklist regions of mm10 obtained from ENCODE were filtered out. BigWig from replicates were merged by first averaging the replicates when converting to a merged bedGraph using wiggleTools:write_bg (1.2.11), then converting back to bigWig by uscs:bedGraphToBigWig (469). Signal tracks were visualized in Interactive Genome Viewer (IGV) (2.16.1) with a consistent data range across all CUT&Tag and ATAC tracks to enable direct comparison. For analysis of rDNA, cleaned reads after fastp removal of adaptors and polyG were mapped to mouse rDNA sequence (NCBI BK000964.3) (Grozdanov et al., 2003) with Bowtie2 using the same parameter. Alignments with MAPQ <30 were filtered out using samtools. Merged rDNA bigWig tracks merged from replicates were generated in the same manner as of above. rDNA signals were visualized in IGV with published curated annotations studying hematopoietic TFs (Antony et al., 2022). Raw and processed data files were deposited in the Gene Expression Omnibus (GEO) under accession number GSE296889.

Peaks and normalized bigWig tracks were used for downstream analysis. Profile and heatmap plots in genebody and around TSS were generated using deepTools. Distance to Ddx21 peak was generated by deepTools using midpoint of Ddx21 peak as “TSS”. Pearson correlation of bigWig among targets was generated using deepTools:plotCorrelation. To obtain co-binding peaks among H3K4me3, Ddx21 and Kdm5a, peak intersection was performed using bedtools:intersect where peaks were intersected if there is 1bp overlap. Peak distribution in the genome was analyzed using CHIPseeker (1.40.0).

For analysis of promoter states, promoters were defined as +/-1500bp around TSS. Combination of presence of histone binding peaks within the promoter and presence of RNA expression (CPM>1) were used to define promoter states: active (promoter H3K4me3+, genebody H3K27me3-, RNA+), poised (promoter H3K4me3+, genebody H3K27me3-, RNA-), bivalent (promoter H3K4me3+, promoter H3K27me3+) and repressed (Promoter H3K4me3-, genebody H3K27me3+). For analysis of enhancers, enhancers were defined as +/- 3000bp regions around H3K4me1 peaks which are located outside of promoters. Combination of presence of histone and ATAC peaks were used to define enhancer states: poised enhancer (H3K27me+, H3K4me1+), primed enhancer (H3K27me3-, H3K4me1+, H3K27Ac^Low^, ATAC^Low^) and active enhancer (H3K27me3-, H3K4me1+, H3K27Ac^Mid/High^, ATAC^Mid/High^) (Barral and Dejardin, 2023).

CUT&Tag signal counts in promoter (+/-1500bp of TSS) were obtained using featureCounts and multiplied with respective scaling factor as stated above. Normalized RNA counts were mapped to promoter counts by matching promoters. Counts were converted to Log_2_ (Counts+1) for spearman correlation using ggpubr (0.6.0). Similarly, counts in Ddx21+ promoter were obtained by counting reads in promoters with Ddx21 peak using featureCounts, and subsequently scaled. Log_2_FC of counts between Ddx21^cKO^ and Ddx21^WT^ were obtained 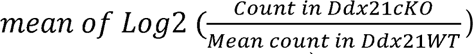. Comparative GO enrichment among Kdm5a FC clusters (Log_2_FC cutoff = 0.2) was performed using clusterProfiler.

Differential binding analysis was performed using Diffbind (3.18.0) with TMM normalization. Consensus peak list was created by only considering peaks that appear in over half of the samples. Differential binding peaks (FC > 0.3, p.adj < 0.05) were mapped to nearest promoter using CHIPpeakAnno (3.8.1). Peaks located within promoters (+/-1500bp of TSS) were used for subsequent GO enrichment analysis.

### ATAC-seq and data analysis

ATAC-seq libraries were constructed in biological duplicates using Hyperactive ATAC-Seq Library Prep Kit for Illumina (Vazyme #TD711) and TruePrep Index Kit V2 for Illumina (Vazyme #TD202) according to product manual. 65000 cells were used per library with 13 cycle of PCR amplification. Single size exclusion was performed to remove small fragments using beads provided in the kit. Libraries were validated by Agilent high sensitivity DNA kit. PE150 sequencing (NovaSeq X) was performed by HaploX.

Downstream processing of ATAC-seq was similar to CUT&Tag with slight differences. Duplicates were removed with Picard and peaks were extended with (--nomodel --shift 100 --ext 200) to account for Tn5 targeting during macs2 peak calling. Raw and processed data files were deposited in the Gene Expression Omnibus (GEO) under accession number GSE296888.

### Statistical analysis

2-tailed unpaired student’s t-test was used to calculate statistical significance in all immunofluorescence, flow cytometry and biochemical analysis. Graphs were plotted using GraphPad Prism 10.

## Acknowledgments

The authors thank the Centre for Regenerative Medicine and Health (CRMH) for providing A.O.L. and C.L. with fellowship, and Prof. Kathy O.L. Lui (Department of Chemical Pathology, CUHK) for insightful discussions.

This work was supported by a grant from Research Grants Council of the Hong Kong Special Administrative Region, China (Project No. 14104321) and a grant from Health@InnoHK Program launched by Innovation Technology Commission of the Hong Kong Special Administrative Region, China.

## Authorship Contributions

Contribution: A.O.L., C.L., and H.H.C designed the study, interpreted the results and wrote the manuscript; A.O.L. and C.L. performed the major experiments; Z.F. generated recombinant DDX21 protein; X.H., J.L., Y.C., H.M.L., F.Z., X.C., and R.N. helped with some other experiments; A.O.L. performed bioinformatics analysis; K.T.L. provided critical comments and revised the manuscript; H.H.C. led and oversaw the project. All authors approved the manuscript for submission.

## Materials availability

RNA sequencing data, ATAC sequencing data, and CUT&Tag sequencing data have been deposited in the Gene Expression Omnibus (GEO) database with accession numbers GSE296783, GSE296888, and GSE296889, respectively. All other data are available upon reasonable request from the corresponding author, Hoi-Hung Cheung.

