## Supplementary Figures and Tables for "RNA helicase Ddx21 safeguards fetal HSPC expansion by recruiting Kdm5a to epigenetically sustain ribosomal and hematopoietic gene transcription"

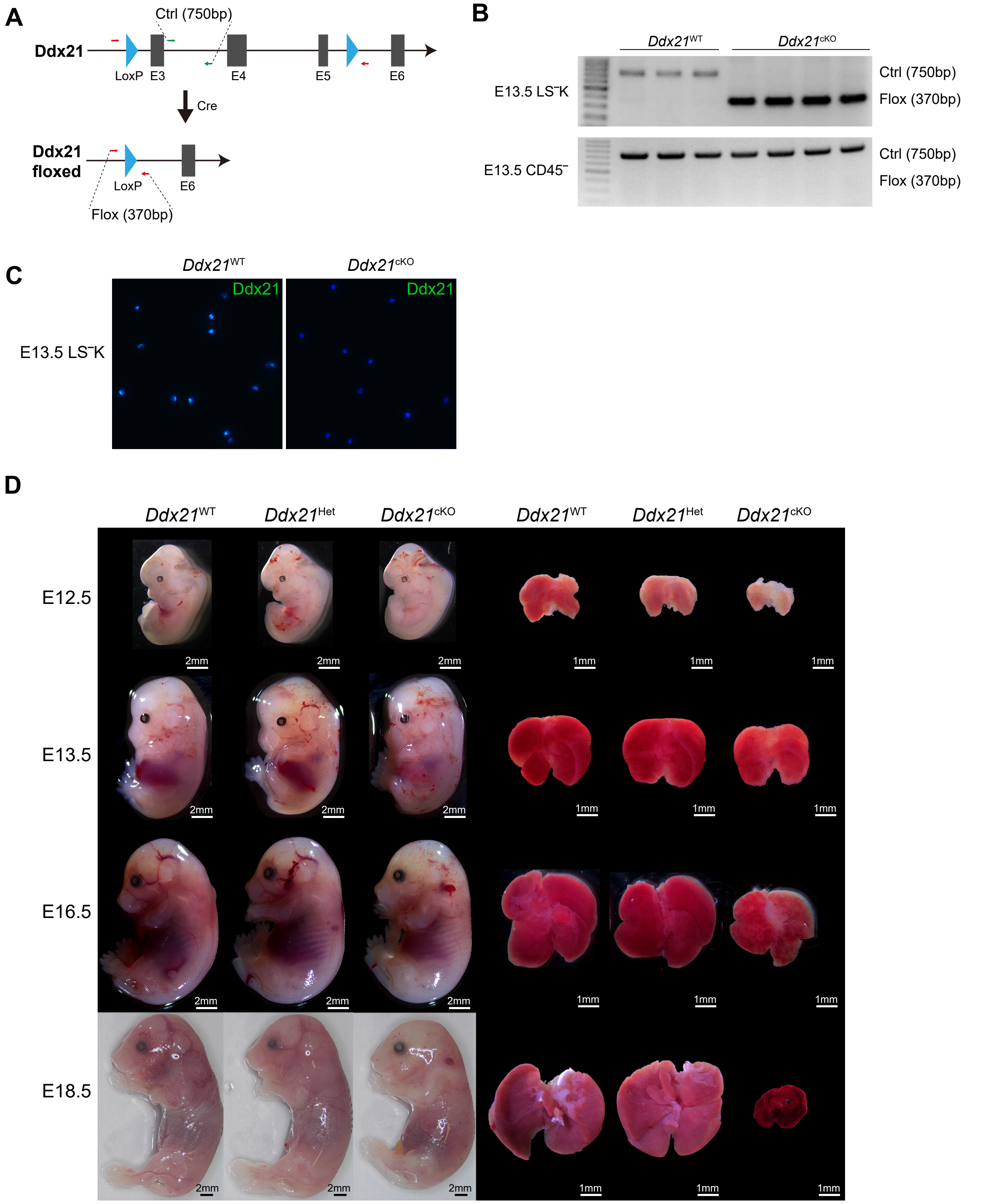


### Supplemental figure 1

(A) Illustration of *Ddx21*$^{flox}$ knockout allele after Cre recombination.

(B) Genomic Ddx21 knockout efficiency by genomic PCR, demonstrating complete and specific knockout of Ddx21 in fetal liver LS$^-$K cells with no deletion in non-hematopoietic cells (CD45$^-$) fetal liver cells of E13.5 *Ddx21*$^{cKO}$ embryos (n=3-4).

(C) Ddx21 immunofluorescence staining of sorted fetal liver LK cells of E13.5 *Ddx21*$^{WT}$, *Ddx21*$^{Het}$ and *Ddx21*$^{cKO}$ embryos.

(D) Gross embryo and fetal liver morphology of E12.5, E13.5, E16.5 and E18.5 *Ddx21*$^{WT}$, *Ddx21*$^{Het}$ and *Ddx21*$^{cKO}$ embryos (n=3-6).

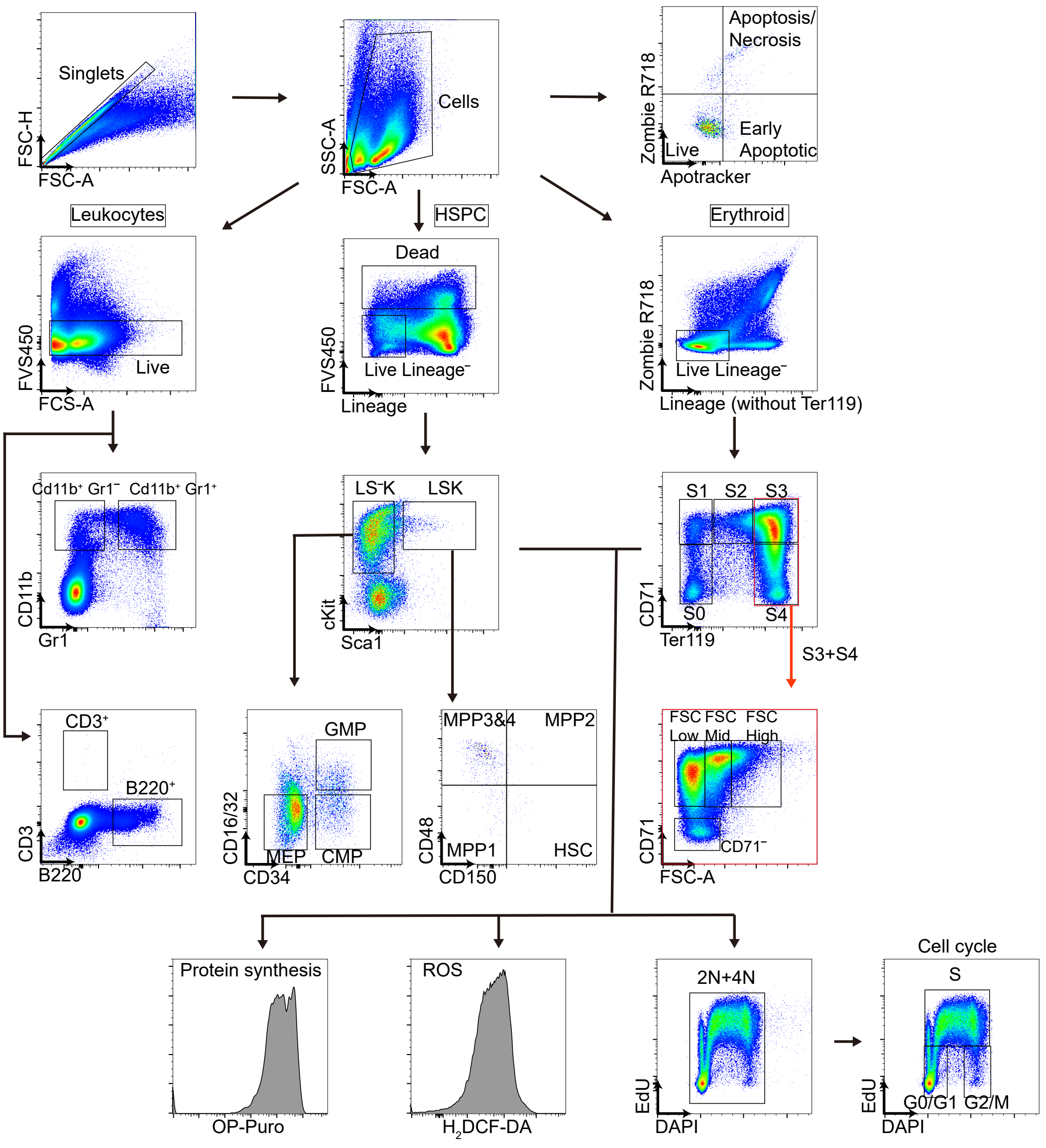


### Supplemental figure 2

Gating strategies for HSPC, erythroids and leukocyte population analysis in fetal liver, along with click-iT alexa fluor 647 detection of 2′,7′-Dichlorodihydrofluorescein diacetate ($H_2$DCF-DA) to detect reactive oxygen species (ROS), O-Propargyl-Puromycin (OP-Puro) to detect nascent protein synthesis and 5-ethynyl-2′-deoxyuridine (EdU) for cell cycle analysis after subsequent staining with 4′,6-diamidino-2-phenylindole (DAPI).

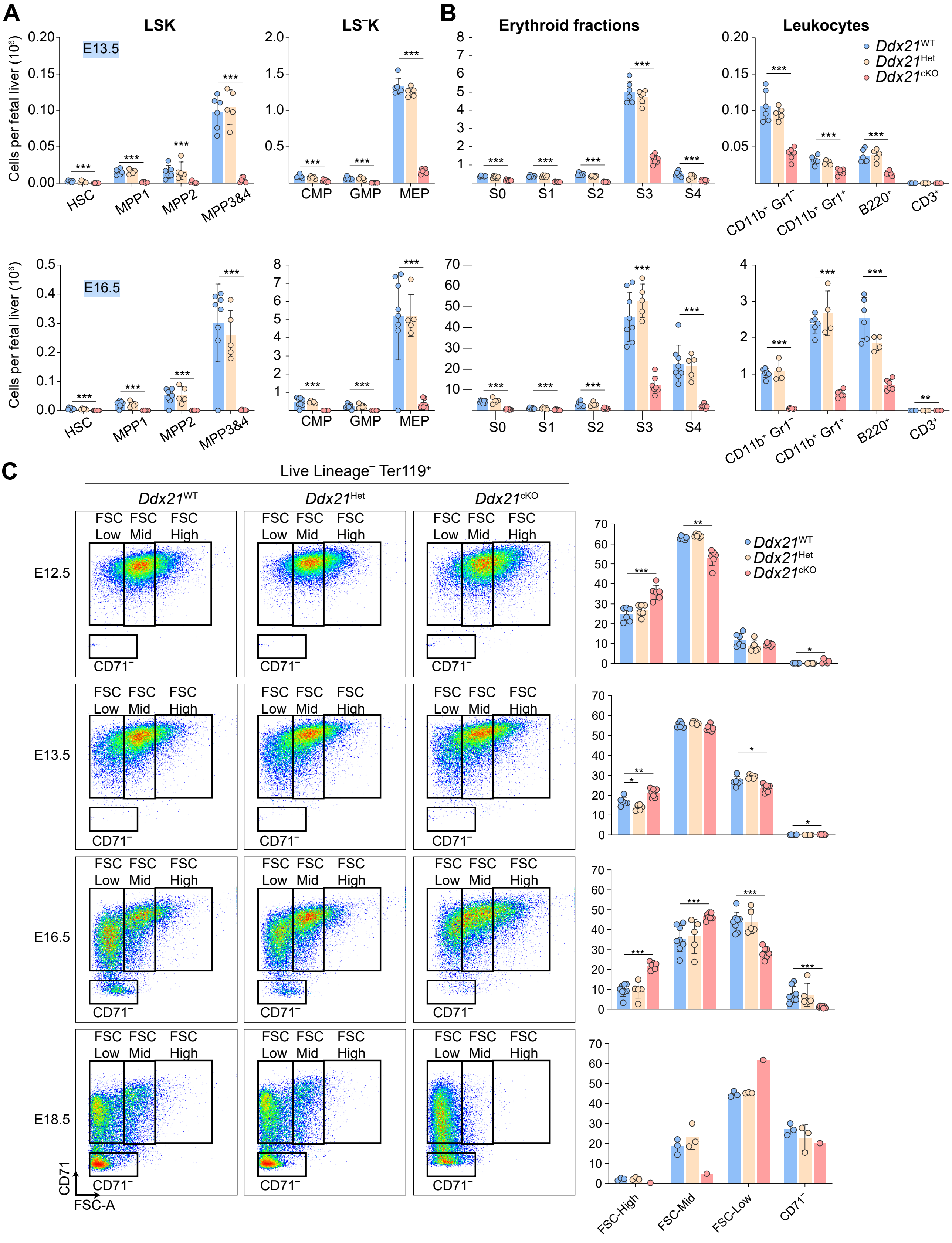


### Supplemental figure 3

(A) Absolute number of hematopoietic stem and progenitor populations of E13.5 and E16.5 *Ddx21*$^{WT}$, *Ddx21*$^{Het}$ and *Ddx21*$^{cKO}$ fetal liver by flow cytometry.

(B) Absolute number of erythroid fractions and leuckotyes of E13.5 and E16.5 *Ddx21*$^{WT}$, *Ddx21*$^{Het}$ and *Ddx21*$^{cKO}$ fetal liver by flow cytometry.

(C) CD71 and FSC flow cytomery analysis of fetal liver terminal erythroid (Live Lineage- Ter119+) differentiation of E12.5 to E18.5 *Ddx21*$^{WT}$, *Ddx21*$^{Het}$ and *Ddx21*$^{cKO}$ embryos (n=1-8). Representative flow plots are shown.

*:<0.05 **:<0.01 ***: <0.001

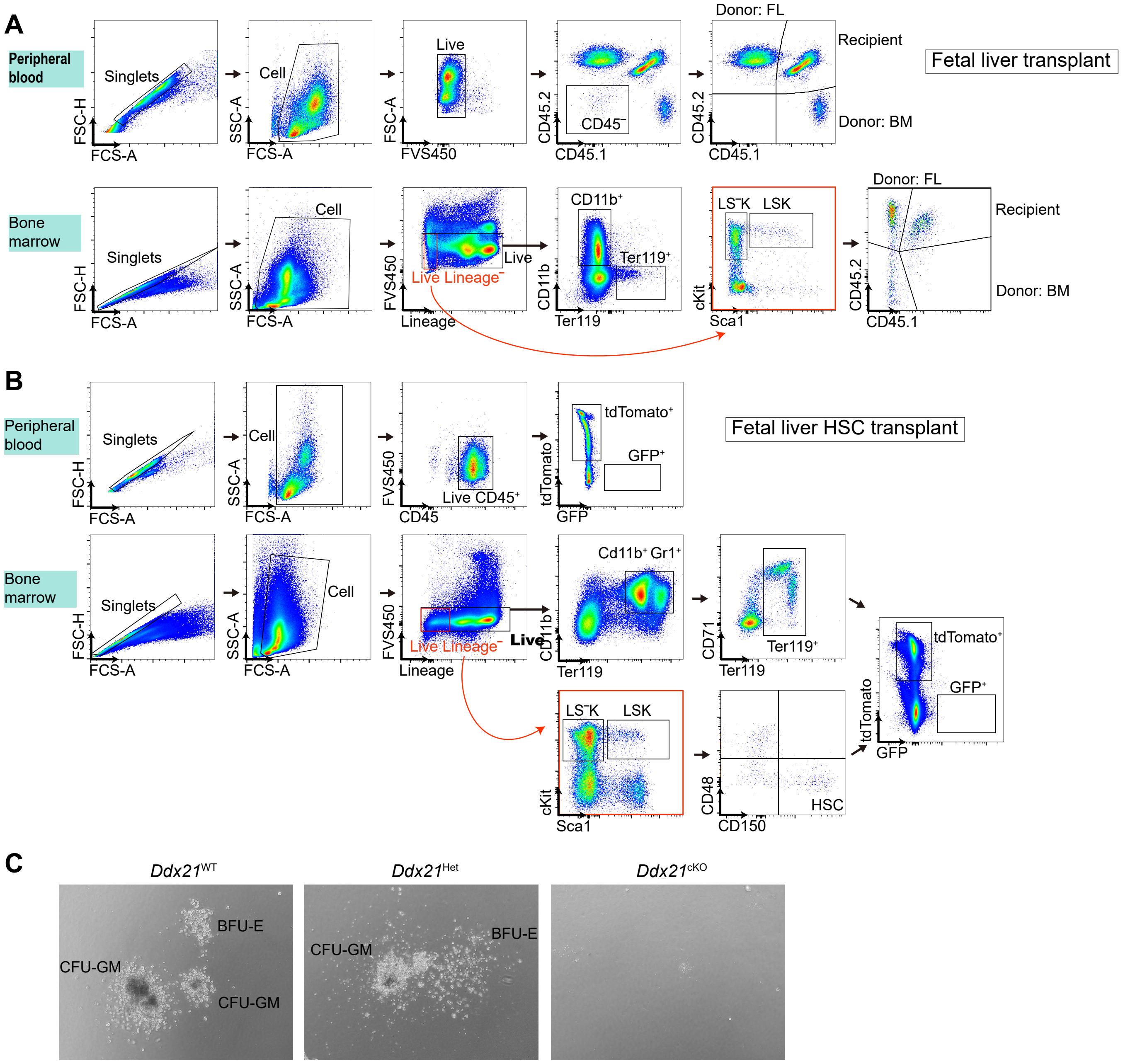


### Supplemental figure 4

(A) Gating strategies of peripheral blood and bone marrow analysis of mice transplanted with whole fetal liver cells.

(B) Gating strategies of peripheral blood and bone marrow analysis of mice transplanted with mTmG$^{f/+}$ fetal liver HSC.

(C) Representative images of colonies generated by CFU assay (methocult M3434) from E13.5 fetal liver cells.

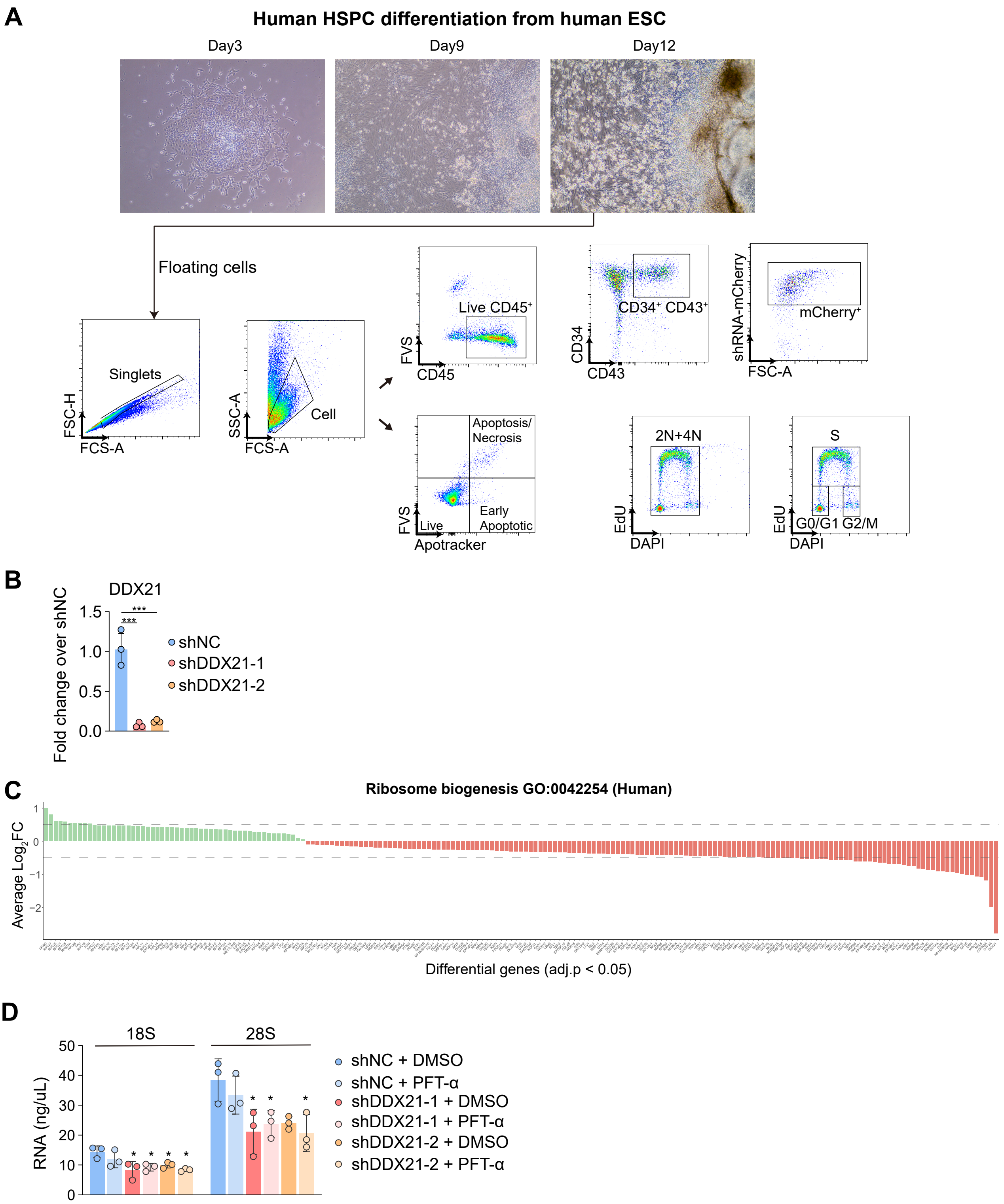


### Supplemental figure 5

(A) Brightfield images and gating strategies of human HSPC differentiation from human ESC.
(B) RT-qPCR knockdown efficiency of DDX21 in human HSPC.
(C) Average $Log_2$ fold change of significant differential genes (adj.p <0.05) between shDDX21 and shNC human HSPC in ribosome biogenesis GO term (0042254).
(D) RNA tapestation analysis of human HSPC after 3 days of shDDX21 lentirivirus transduction and 15μM Pifithrin-α (PFT-α) incubation.

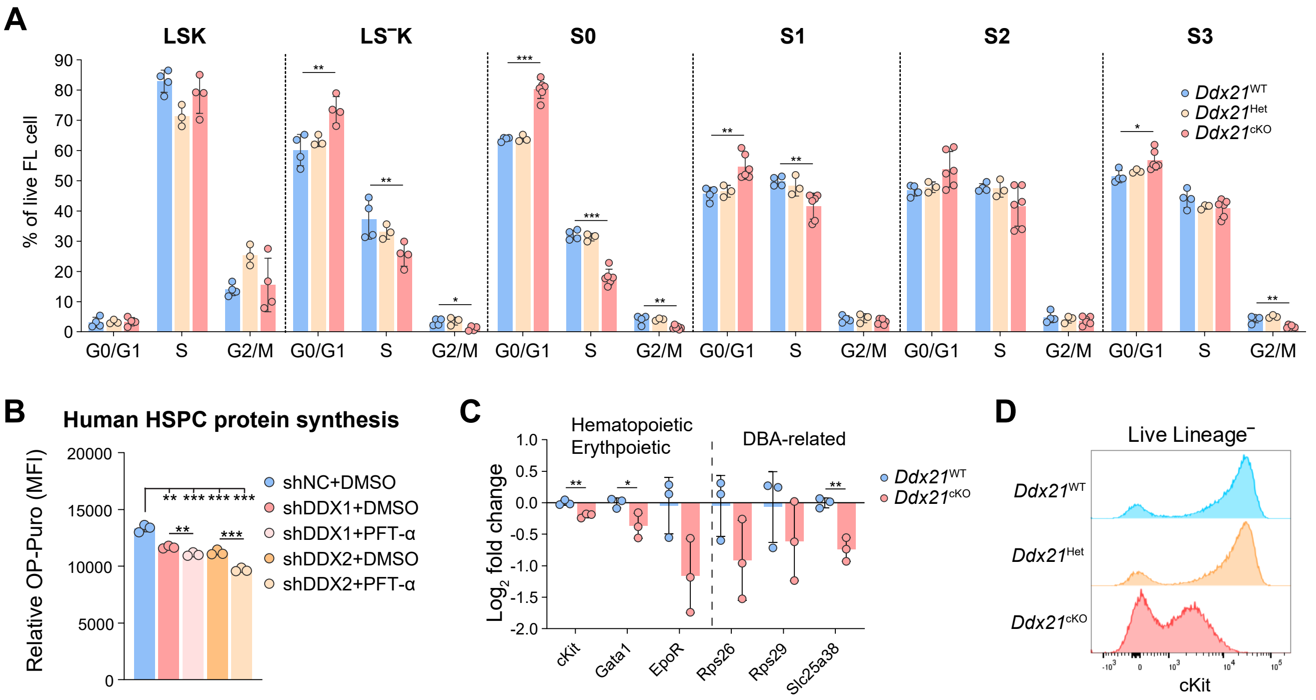


### Supplemental figure 6

(A) EdU/DAPI cell cycle analysis of E13.5 fetal liver cells 1.5hr post EdU injection into pregnant mice.
(B) OP-Puro geometric mean fluorescence of human HSPC after 2 days of knockdown and culture in 15μM Pifithrin-α (PFT-α).
(C) Log2 fold change of erythropoietic genes and Diamond Blackfan Anemia (DBA) genes in mouse HPSC transcriptome.
(D) Flow plot of cKit fluorescence in Live Lineage⁻ cells of E13.5 fetal liver.
*:<0.05 **:<0.01 ***: <0.001

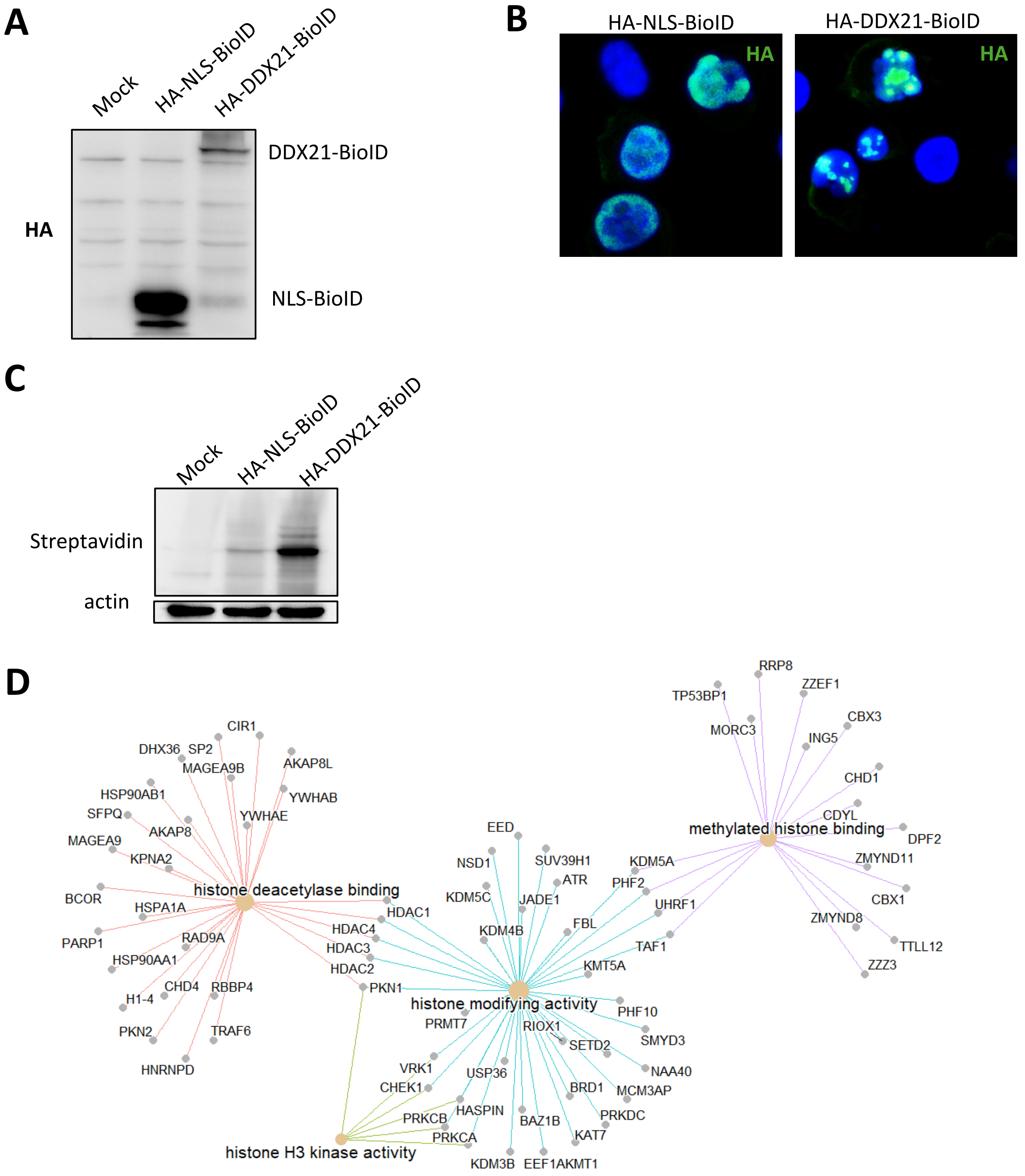


**Supplemental figure 7**

(A-E) DDX21-BioID2 interactome analysis. K562 cells were transduced with either HA-NLS-BioID or HA-DDX21-BioID lentivirus. Cells were expanded for 5 days before biotin was added. Biotinylated protein were subjected to LC-MS/MS (4D-DIA).

(A) HA western blot showing expression of BioID fusion proteins.
(B) HA immunofluorescence showing nuclear localization of BioID fusion proteins.
(C) Streptavidin western blot of whole cell lysate after 1 day of biotin incubation.
(D) Molecular function GO terms related to histone modifications were enriched in the DDX21 interactome.

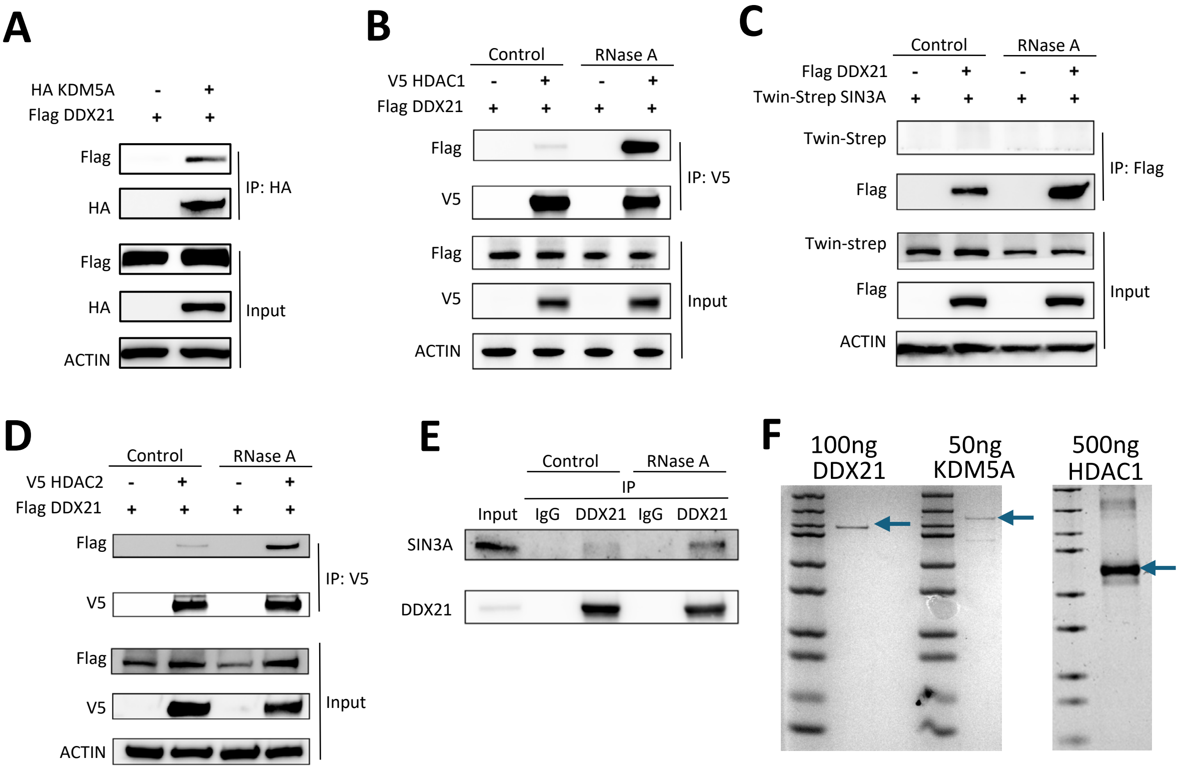


### Supplemental figure 8

(A) Ectopic KDM5A co-immunoprecipitation with ectopic DDX21 in 293F

(B) Ectopic HDAC1 co-immunoprecipitation with ectopic DDX21 in 293F with or without RNase A treatment.

(C) Ectopic DDX21 co-immunoprecipitation with ectopic SIN3A in 293F with or without RNase A treatment.

(D) Ectopic HDAC2 co-immunoprecipitation with ectopic DDX21 in 293F with or without RNase A treatment.

(E) Endogenous DDX21 co-immunoprecipitation with endogenous SIN3A in K562 with or without RNase A treatment.

(F) Coomassie blue staining of purified DDX21, KDM5A and HDAC1 proteins.

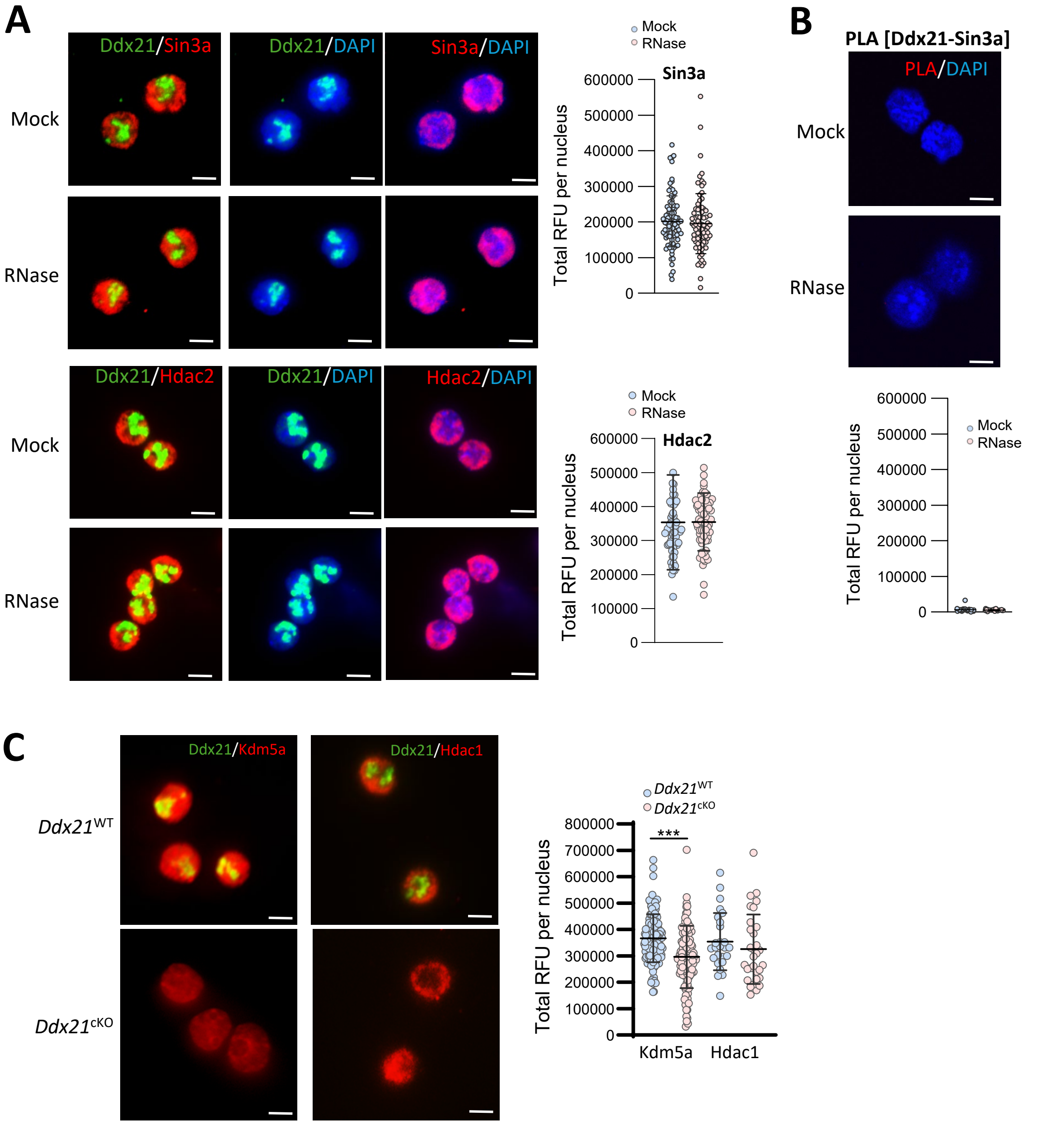


### Supplemental figure 9

(A) Immunofluorescence of Ddx21, Sin3a and Hdac2 in Mock and RNase A treated E13.5 fetal liver LK cells. Total relative fluorescence within the nucleus was quantified. Scale bar = 5µm.

(B) Proximity ligation assay (PLA) between Ddx21 and Sin3a in mock and RNase A treated E13.5 fetal liver LK cells. Total relative fluorescence within the nucleus was quantified. Scale bar = 5µm.

(C) Ddx21, Kdm5a and Hdac1 immunofluorescence in *Ddx21*$^{WT}$ and *Ddx21*$^{cKO}$ E13.5 fetal liver LK cells. Total relative fluorescence within the nucleus was quantified. Scale bar = 5µm.

*:<0.05 **:<0.01 ***: <0.001

Ddx21-Kdm5a associate to active promoter and occasionally to TES but not repressive mark

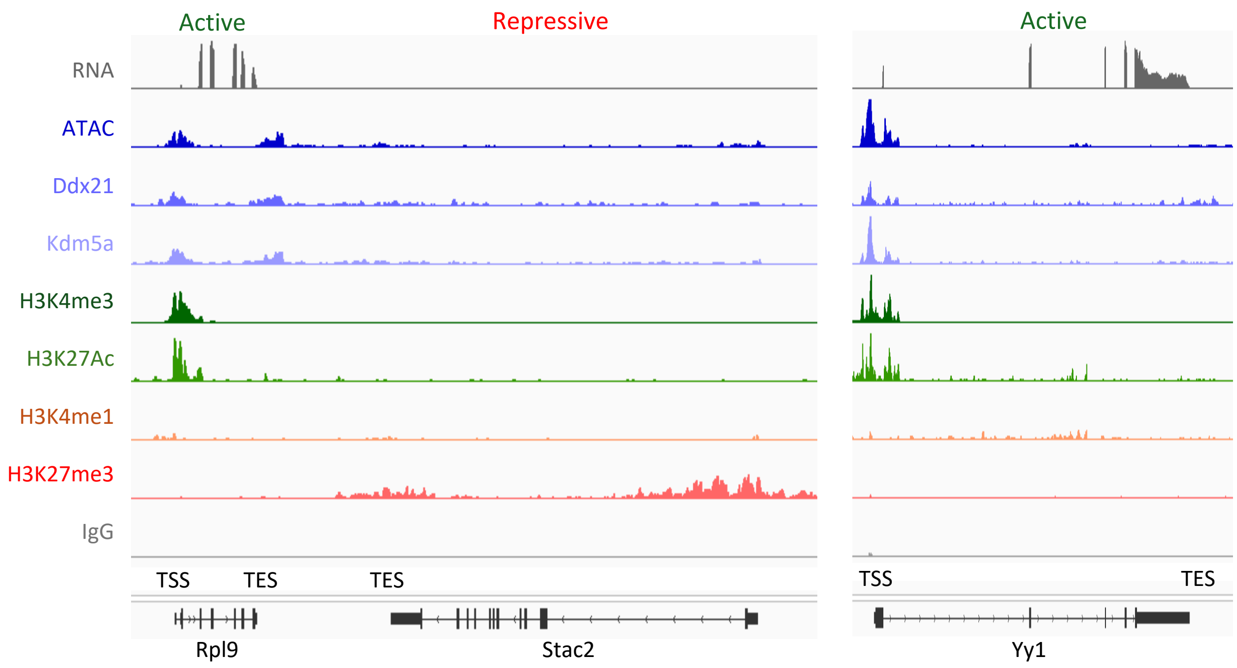


Ddx21 and Kdm5a associates to enhancer marks and does not bind to all active promoters

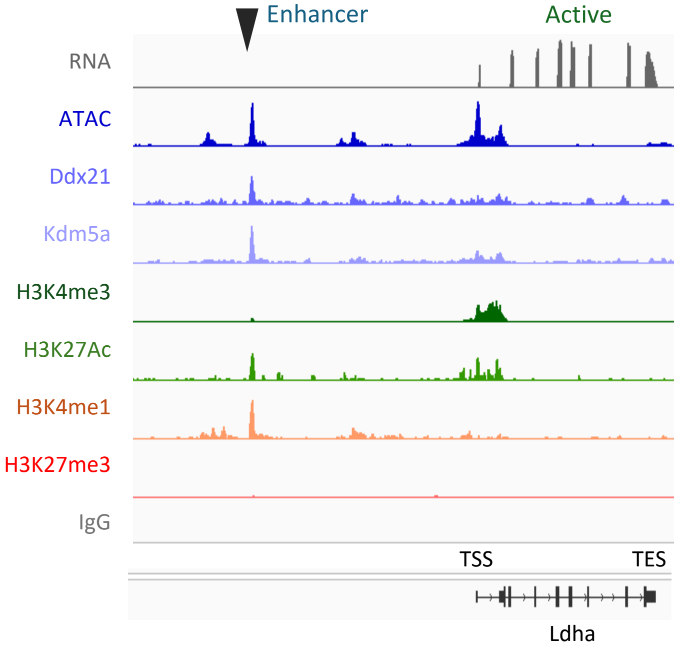


**Supplemental figure 10**

Representative tracks of RNA level, ATAC and CUT&Tag signals of fetal liver LK cells from E13.5 *Ddx21*$^{WT}$ embryos, demonstrating Kdm5a and Ddx21 binding in active promoter and enhancer but not repressive promoter. TSS: Transcription start site. TES: Transcription end site.

### Ooccupancy of histone marks, ATAC, Kdm5a and Ddx21 around TSS

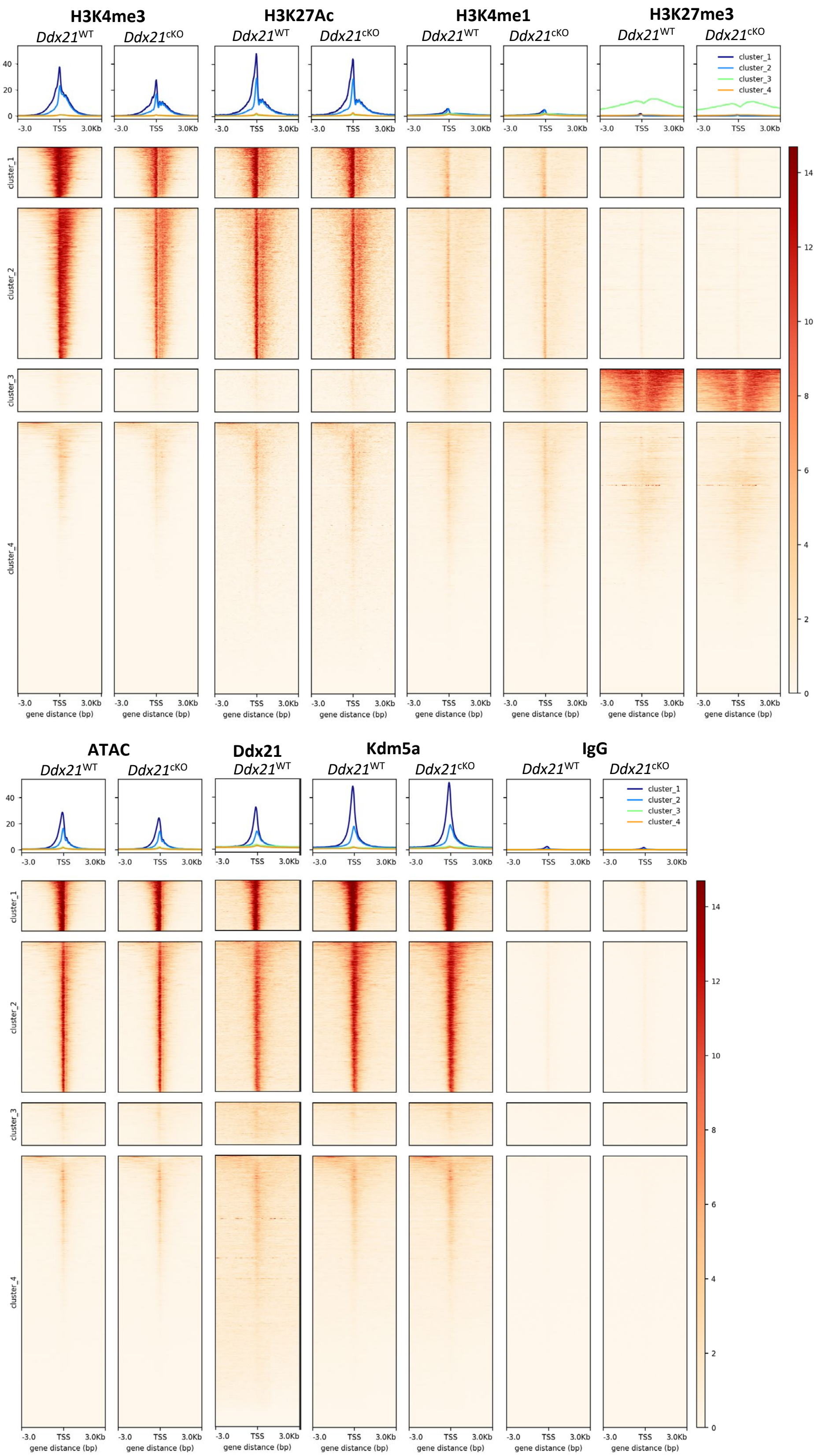


### Supplemental figure 11

Profile and heatmap plot of TMM-background bin normalized and per-million scaled CUT&Tag signals of H3K4me3, H3K4me1, H3K27me3, H3K27Ac, ATAC, Kdm5a, Ddx21 and IgG around TSS of E13.5 *Ddx21*$^{WT}$ and *Ddx21*$^{cKO}$ fetal liver LK cells. TSS: Transcription start site.

### Ooccupancy of histone marks, ATAC, Kdm5a and Ddx21 in gene body

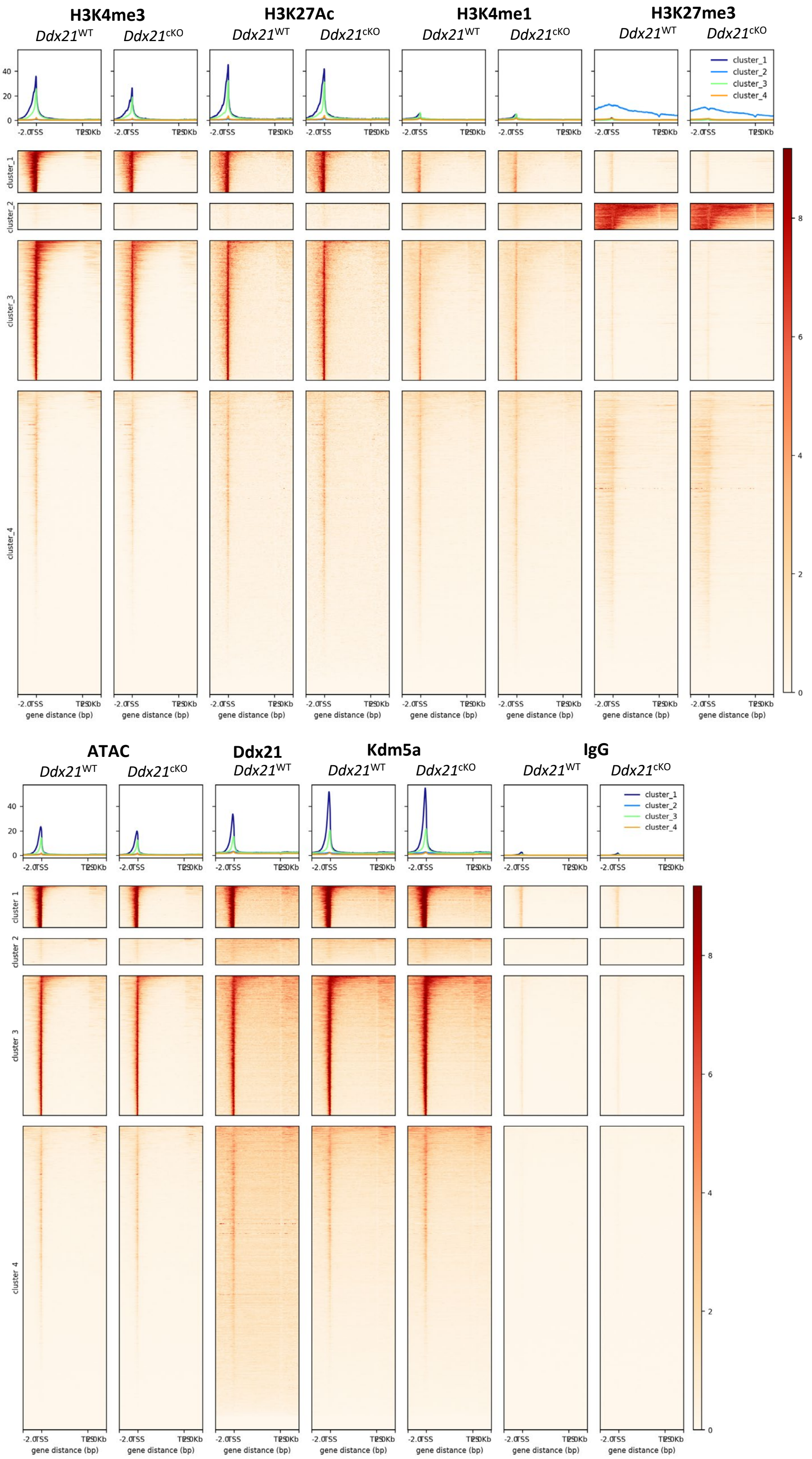


### Supplemental figure 12

Profile and heatmap plot of TMM-background bin normalized and per-million scaled CUT&Tag signals of H3K4me3, H3K4me1, H3K27me3, H3K27Ac, ATAC, Kdm5a, Ddx21 and IgG in gene body of E13.5 *Ddx21*$^{WT}$ and *Ddx21*$^{cKO}$ fetal liver LK cells. TSS: Transcription start site. TES: Transcription end site

### Ooccupancy of histone marks, ATAC, Kdm5a and Ddx21 in H3K4me1+ enhancers

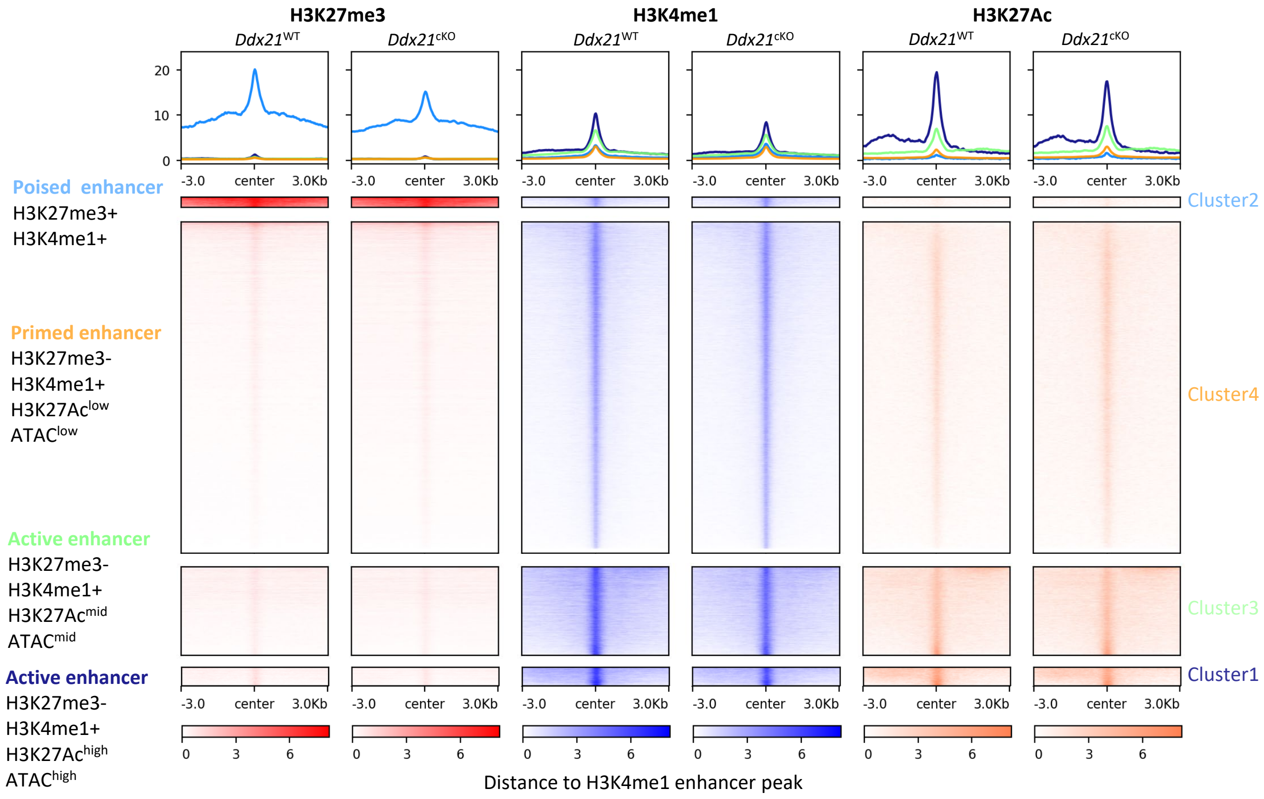


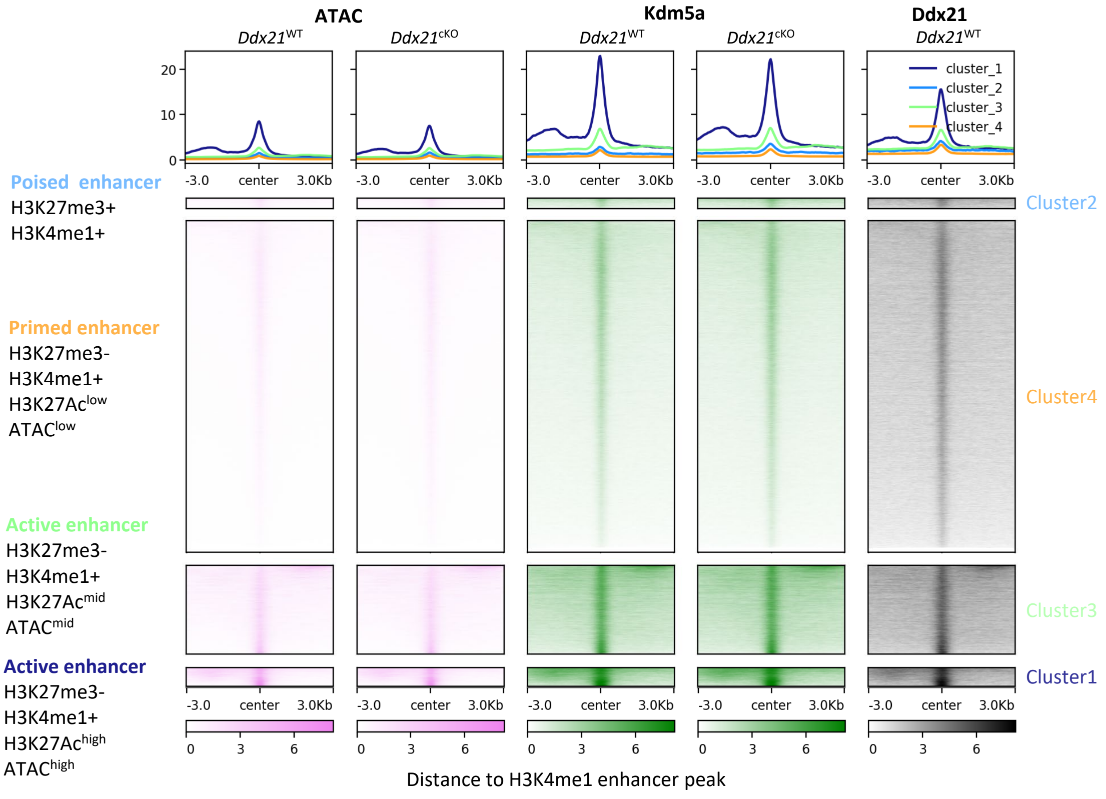


### Supplementary figure 13

Profile and heatmap plot of TMM-background bin normalized and per-million scaled CUT&Tag and ATAC signals in H3K4me1+ enhancer peaks (H3K4me1+ peaks in promoter were excluded). Enhancers were grouped to poised, primed and active enhancers based on combinatorial histone and ATAC binding (Barral A et al., 2023).

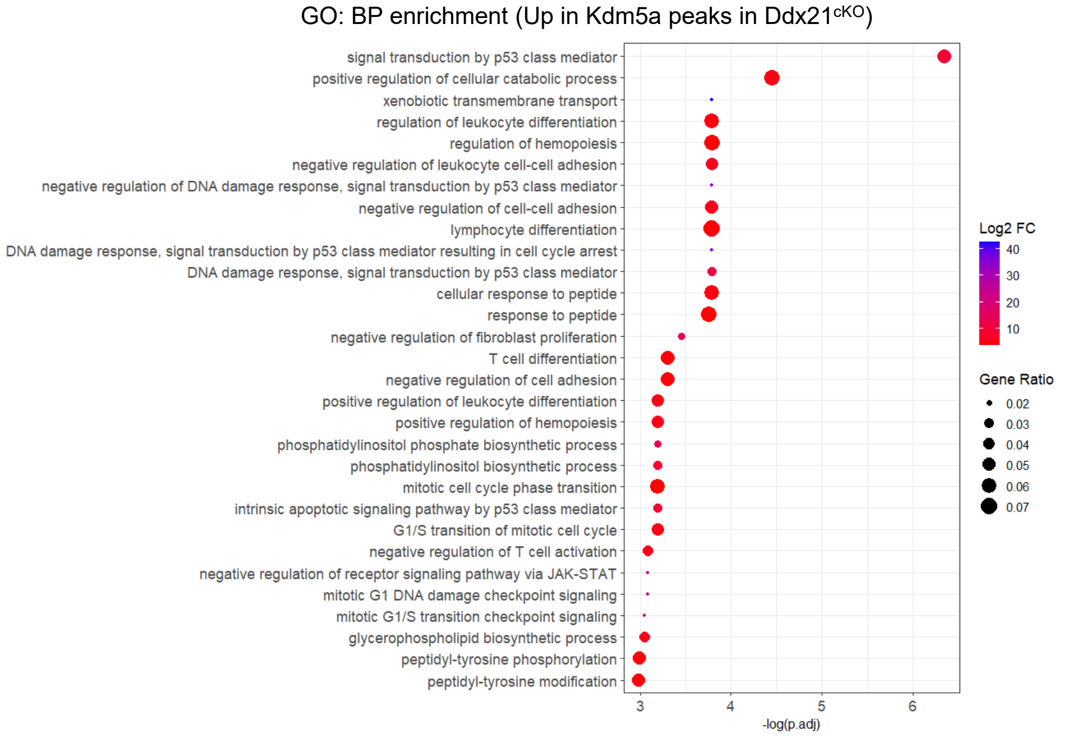


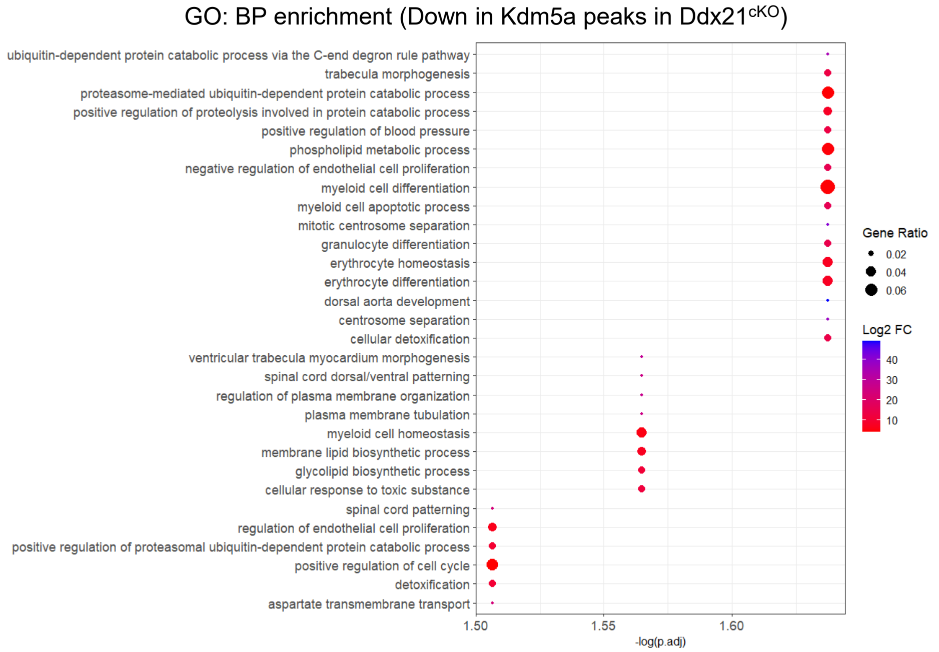


### Supplemental figure 14

Biological processes GO enrichment of genes with decrease or increase in CUT&Tag Kdm5a binding in *Ddx21*$^{cKO}$ compared to *Ddx21*$^{WT}$ E13.5 fetal liver LK cells using Diffbind with TMM normalization.

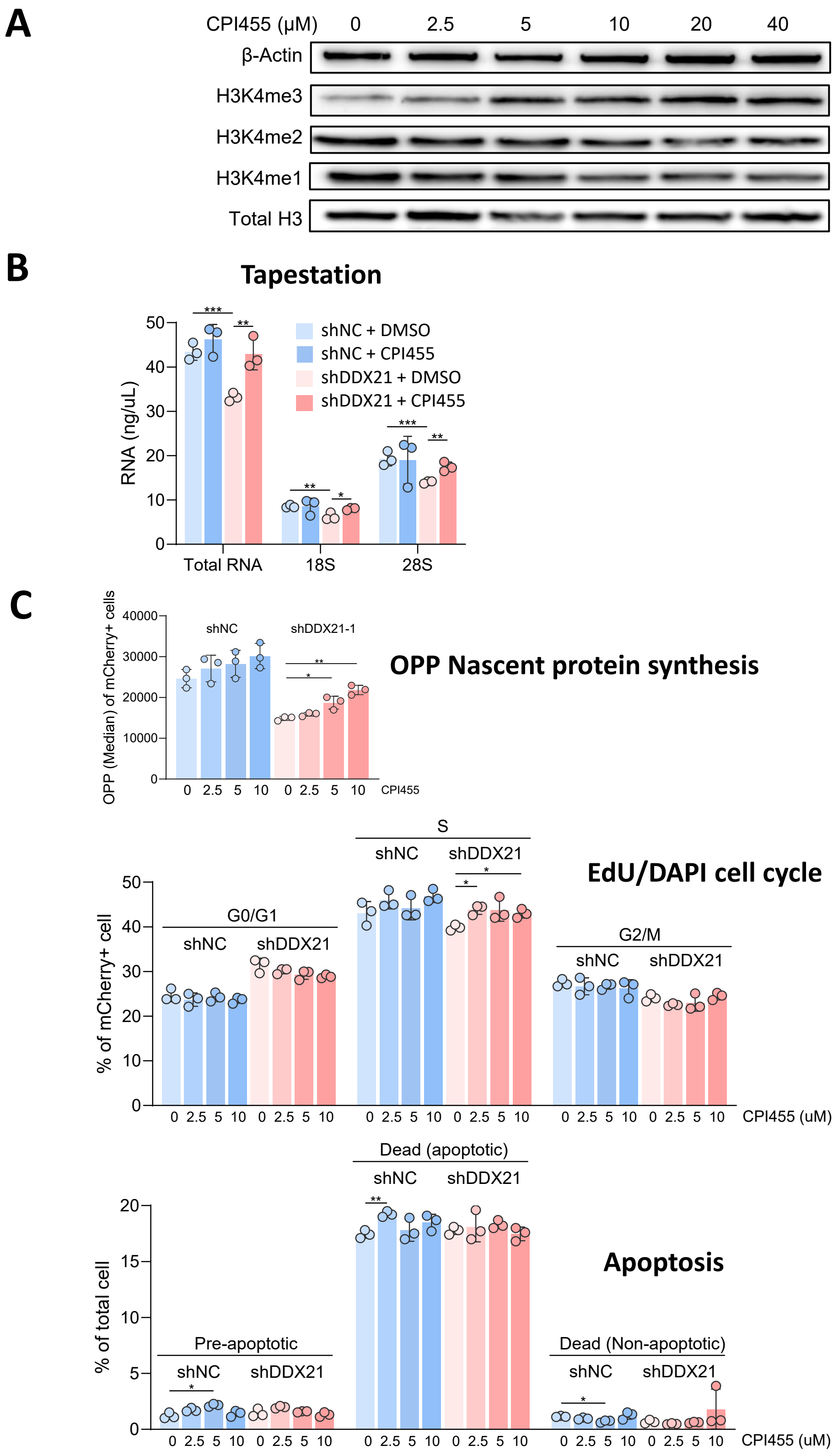


### Supplemental figure 15

(A) Western blot of H3K4me3, H3k4me2, H3K4me1, total H3 and β-Actin of K562 cells after 1 day of CPI455 treatment.

(B) TapeStation rRNA analysis of K562 cells 2 days post transduction with shRNA lentivirus and culture in 10μM CPI455/DMSO (n=3).

(C) OP-Purio nascent protein synthesis, EdU/DAPI cell cycle and apoptosis/cell death flow cytometry analysis of K562 cells 2 days post lentiviral transduction and culture with CPI455/DMSO. OP-Puro was incorporated for 30min, EdU for 1hr.

*: p<0.05, **: p<0.01, ***: p<0.001

A

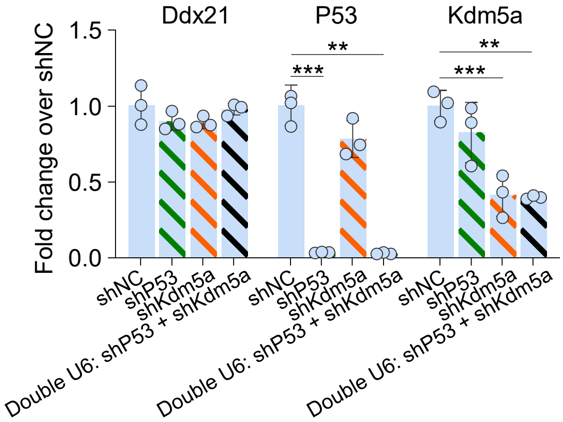


### Supplemental figure 16

(A) RT-qPCR analysis of shRNA knockdown efficiency in cultured E13.5 LK cells after 3 days of lentiviral transduction carrying either single or double U6-shRNA construct (n=3).
*:<0.05 **:<0.01 ***: <0.001

| Purpose | F | R | Tm | Size (bp) |
|---|---|---|---|---|
| Vav1-Cre genotyping | GGTGTTGTAGTTGTCCCCACT | CAGGTTTTGGTGCACAGTCA | 60 | Cre: 390<br>WT: 212 |
| Ddx21$^{flox}$ genotyping | GAACTCAGAGATCTGCCTCTCAAAT | CCGAGTGTTCTCTTGTGCTGTA | 60 | flox: 201<br>WT: 144 |
| mTmG$^{flox}$ genotyping | AGGGAGCTGCAGTGGAGTAG | CTTTAAGCCTGCCCAGAAGA | 60 | flox: 128<br>WT: 212 |
| | TAGAGCTTGCGGAACCCTTC | | | |
| Ddx21$^{flox}$ KO efficiency | GAACTCAGAGATCTGCCTCTCAAAT | CATAAGAAAGTGAGAGGGCACAGA | 63 | Delta: 370 |
| | GTATCAGGGAGCAAACCCCA | TTGTGCGGTTTGAGGTCAGT | 63 | Flox: 750 |

### Supplemental table 1

Primers used for mouse genotyping and Ddx21$^{flox}$ KO efficiency test

| Name | Function | Stem | Loop |
|---|---|---|---|
| shNC | shRNA control | CCTAAGGTTAAGTCGCCCTCG | CCTGACCCA |
| shDdx21 | Mouse Ddx21 KD | GTCTCCAGACAAGGGTATATA | CCTGACCCA |
| shP53 | Mouse P53 KD | GTACATGTGTAATAGCTCC | CCTGACCCA |
| shKdm5a | Mouse Kdm5a KD | CGAGGAAATAATGAGGATAAA | CCTGACCCA |
| shDDX21-1 | Human DDX21 KD | CACCGCCCATATCTGAAGAAACTATT | CCTGACCCA |
| shDDX21-2 | Human DDX21 KD | CACCGCATGAGGAATGGGATTG | ATACTCGAGTAT |
| shHuP53 | Human P53 KD | GACTCCAGTGGTAATCTAC | CCTGACCCA |

### Supplemental table 2

Stem and loop sequences of shRNAs

| **Lineage-biotin panel for fetal liver population analysis**<br>For erythroid fraction analysis, Ter119-biotin is omitted and zombie-R718 is used | | | | |
|---|---|---|---|---|
| **Target** | **Conjugate** | **Cat no.** | **Manufacturer** | **Dilution** |
| CD3 | Biotin | 100303 | Biolegend | 1:125 |
| B220 | Biotin | 103203 | Biolegend | 1:125 |
| Gr1 | Biotin | 108403 | Biolegend | 1:125 |
| CD11b | Biotin | 101203 | Biolegend | 1:125 |
| Ter119 | Biotin | 116203 | Biolegend | 1:125 |
| Dead (HSC/leukocyte) | FVS450 | 562247 | BD | 1:250 |
| Dead (erythroid) | Zombie-R718 | 423115 | Biolegend | 1:250 |

| **Fetal liver HSPC surface panel** | | | | |
|---|---|---|---|---|
| **Target** | **Conjugate** | **Cat no.** | **Manufacturer** | **Dilution** |
| Biotin | Streptavidin-APC/Cy7 | 405208 | Biolegend | 1:125 |
| cKit | APC | 135107 | Biolegend | 1:125 |
| Sca1 | BV605 | 108133 | Biolegend | 1:125 |
| CD48 | PE | 103405 | Biolegend | 1:125 |
| CD150 | BV510 | 115929 | Biolegend | 1:125 |
| CD16/32 | PerCP/Cy5.5 | 101323 | Biolegend | 1:125 |
| CD34 | FITC | 11-0341-81 | Life Technologies | 1:125 |

| **Fetal liver erythroid fraction surface panel** | | | | |
|---|---|---|---|---|
| **Target** | **Conjugate / dye** | **Cat no.** | **Manufacturer** | **Dilution** |
| Biotin (lineage) | Streptavidin-APC/Cy7 | 405208 | Biolegend | 1:125 |
| CD71 | BV510 | 113823 | Biolegend | 1:125 |
| Ter119 | PE/Cy7 | 116221 | Biolegend | 1:125 |
| DNA | Hoechst33342 | H1399 | Life Technologies | 1ug/ml |
| ROS | $H_2$DCF-DA | D399 | Life Technologies | 10µM |

| **Fetal liver leukocytes surface panel** | | | | |
|---|---|---|---|---|
| **Target** | **Conjugate** | **Cat no.** | **Manufacturer** | **Dilution** |
| CD3 | PE | 100307 | Biolegend | 1:125 |
| B220 | APC/Cy7 | 103223 | Biolegend | 1:125 |
| Gr1 | APC | 108411 | Biolegend | 1:125 |
| CD11b | FITC | 101205 | Biolegend | 1:125 |

| **Apoptosis** | | | | |
|---|---|---|---|---|
| **Target** | **Conjugate** | **Cat no.** | **Manufacturer** | **Dilution** |
| Apoptotic cell | Apotracker Green | 427402 | Biolegend | 1:125 |
| Dead | FVS450 | 562247 | BD | 1:250 |

| **Cell cycle Click-iT (After surface stain, fix, permeabilize)** | | | | |
|---|---|---|---|---|
| **Target** | **Conjugate** | **Cat no.** | **Manufacturer** | **Dilution** |
| EdU | Alexa fluor 647 | C10419 | Life Technologies | As in manual |
| DNA | DAPI | D1306 | Life Technologies | 0.1ug/ml |

| **Nascent protein synthesis Click-iT (After surface stain, fix, permeabilize)** | | | | |
|---|---|---|---|---|
| **Target** | **Conjugate** | **Cat no.** | **Manufacturer** | **Dilution** |
| OP-Puro | Alexa fluor 647 | C10458 | Life Technologies | As in manual |

### Supplemental table 3

All antibodies, reagents and dyes used in flow cytometry and sorting for fetal liver population analysis, apoptosis and cell cycle.

| CD45.1/CD45.2 transplantation analysis and HSC sorting for transplantation | | | | | |
|---|---|---|---|---|---|
| **Target** | **Conjugate** | **Cat no.** | **Manufacturer** | **Dilution** | **HSC sorting** |
| **Common** | | | | | |
| CD45.1 | PE/Cy7 | 110729 | Biolegend | 1:125 | |
| CD45.2 | FITC | 109805 | Biolegend | 1:125 | |
| Dead | FVS450 | 562247 | BD | 1:250 | |
| **Peripheral blood** | | | | | |
| CD3 | PerCP | 100325 | Biolegend | 1:125 | |
| B220 | APC/Cy7 | 103223 | Biolegend | 1:125 | |
| CD11b | Spark UV387 | 101291 | Biolegend | 1:125 | |
| **Bone marrow HSPC + Ter119 + Cd11b (Stained with lineage-biotin panel first)** | | | | | |
| cKit | APC | 135107 | Biolegend | 1:125 | * |
| Sca1 | BV605 | 108133 | Biolegend | 1:125 | * |
| CD48 | PE | 103405 | Biolegend | 1:125 | * |
| CD150 | BV510 | 115929 | Biolegend | 1:125 | * |
| Ter119 | PerCP | 116225 | Biolegend | 1:125 | |
| CD11b | SparkUV387 | 101291 | Biolegend | 1:125 | |
| Biotin (lineage) | Streptavidin-APC/Cy7 | 405208 | Biolegend | 1:125 | * |

| $mTmG^{flox}$ transplantation analysis | | | | |
|---|---|---|---|---|
| **Target** | **Conjugate** | **Cat no.** | **Manufacturer** | **Dilution** |
| **Common** | | | | |
| | GFP | | | |
| | Tdtomato | | | |
| Dead | FVS450 | 562247 | BD | 1:250 |
| **Peripheral blood** | | | | |
| CD45 | APC | 103111 | Biolegend | 1:125 |
| CD3 | PerCP | 100325 | Biolegend | 1:125 |
| B220 | APC/Cy7 | 103223 | Biolegend | 1:125 |
| CD11b | Spark UV387 | 101291 | Biolegend | 1:125 |
| **Bone marrow HSPC (Stained with lineage-biotin panel first)** | | | | |
| cKit | APC | 135107 | Biolegend | 1:125 |
| Sca1 | BV605 | 108133 | Biolegend | 1:125 |
| CD16/32 | PerCP/Cy5.5 | 101323 | Biolegend | 1:125 |
| CD34 | AF700 | 560518 | BD | 1:125 |
| CD48 | PE/Cy7 | 103423 | Biolegend | 1:125 |
| CD150 | BV510 | 115929 | Biolegend | 1:125 |
| Biotin (lineage) | Streptavidin-APC/Cy7 | 405208 | Biolegend | 1:125 |
| **Bone marrow erythroid fractions and myeloids (Stained with lineage-biotin panel first)** | | | | |
| CD71 | BV510 | 113823 | Biolegend | 1:125 |
| Ter119 | PE/Cy7 | 116221 | Biolegend | 1:125 |
| Gr1 | APC | 108411 | Biolegend | 1:125 |
| CD11b | SparkUV387 | 101291 | Biolegend | 1:125 |
| Biotin | Streptavidin-APC/Cy7 | 405208 | Biolegend | 1:125 |

| Human HSPC panel | | | | |
|---|---|---|---|---|
| **Target** | **Conjugate** | **Cat no.** | **Manufacturer** | **Dilution** |
| | shRNA-mCherry | | | |
| CD34 | APC-Cy7 | 343513 | Biolegend | 1:125 |
| CD45 | PE | 304007 | Biolegend | 1:125 |
| CD43 | PE-Cy7 | 343207 | Biolegend | 1:125 |
| Dead | FVS450 | 562247 | BD | 1:250 |

### Supplemental table 4

All antibodies, reagents and fluorochromes used in flow cytometry and sorting for mouse transplantation analysis and human HSPC analysis.

| Antibody/Beads/Dyes | Cat no. | Manufacturer | Purpose |
|---|---|---|---|
| **Primary Ab** | | | |
| Rb anti-Ddx21 | Nil (raised) | ABclonal | IF, IP, CUT&Tag, PLA |
| Rb anti-DDX21 | 10528-1-AP | Proteintech | IP, WB |
| Ms anti-Sin3a/SIN3A | 5299 | Santa Cruz | IF, IP, PLA, WB |
| Ms anti-Hdac1/HDAC1 | sc81598 | Santa Cruz | IF, IP, PLA, WB |
| Ms anti-Hdac2/HDAC2 | 5133 | Cell Signaling Technologies | IF, IP, WB |
| Rb anti-Kdm5a/KDM5A | 3876 | Cell Signaling Technologies | CUT&Tag, IP, WB |
| Ms anti-Kdm5a/KDM5A | GTX00688 | Genetex | IF, IP, PLA, WB |
| Rb anti-H3K27Ac | ab4729 | Abcam | CUT&Tag |
| Rb anti-H3K27me3 | 9733 | Cell Signaling Technologies | CUT&Tag |
| Rb anti-IgG | 12-370 | Milipore | CUT&Tag |
| Rb anti-H3K4me3 | 9751 | Cell Signaling Technologies | CUT&Tag, WB |
| Rb anti-H3K4me2 | 9725 | Cell Signaling Technologies | WB |
| Rb anti-H3K4me1 | ab8895 | Abcam | CUT&Tag, WB |
| Rb anti-Total H3 | A2348 | ABclonal | WB |
| Rb anti-β-Actin/ACTIN | 4970 | Cell Signaling Technologies | WB |
| Ms anti-Flag | F1804 | Sigma | WB |
| Rb anti-Flag | PA1-984B | Life Technologies | WB |
| Rb anti-HA | 71-5500 | Life Technologies | IF, WB |
| Rb anti-V5 | 14440-1-AP | Proteintech | IP, WB |
| **Secondary Ab** | | | |
| Goat anti-mouse IgG AF488 | A32723 | Life Technologies | IF |
| Goat anti-rabbit IgG AF647 | A32733 | Life Technologies | IF |
| Goat anti-mouse IgG HRP | 1706516 | Bio-rad | WB |
| Goat anti-rabbit IgG HRP | 1706515 | Bio-rad | WB |
| **Beads** | | | |
| Anti-Flag magnetic beads | M8823 | Sigma | IP |
| Anti-HA magnetic beads | 88836 | Life Technologies | IP |
| Anti-Twin Strep-tag Streptactin-XT magnetic beads | KTSM1350 | AlpaLifeBio | IP |
| Anti-Ig Dynabeads Protein G | 10004D | Life Technologies | IP |
| **Dyes** | | | |
| HRP dye for Twin Strep-tag detection | KTSM1550 | AlpaLifeBio | WB |
| Streptavidin-HRP dye for biotin detection | S2438 | Sigma | WB |

### Supplemental table 5

Antibodies, beads and dyes used in immunofluorescence (IF), immunoprecipitation (IP), CUT&Tag and western blot (WB), and proximity ligation assay staining (PLA).

| Gene | Organism | F | R |
|---|---|---|---|
| Pre-rRNA | Mouse | TTTTGGGGAGGTGGAGAGTC | AGAGAACTCCGGAGCACCAC |
| 18S rRNA | Mouse | AGTCCCTGCCCTTTGTACACA | CGATCCGAGGGCCTCACTA |
| 28S rRNA | Mouse | TGGGTTTTAAGCAGGAGGTG | GTGAATTCTGCTTCACAATG |
| Ddx21 | Mouse | GACTTAATTGCCCAGGCACG | TGCAAGGACTAGCACCTGA |
| P21 | Mouse | AACATCTCAGGGCCGAAAA | TGCGCTTGGAGTGATAGAAA |
| Mdm2 | Mouse | GAAAAGCCTGAGGCTGGTAGAA | AACATAGGCAACCACCAGGAA |
| Bax | Mouse | GGAGCAGCTTGGGAGCG | AAAAGGCCCCTGTCTTCATGA |
| Gapdh | Mouse | TGGCCTTCCGTGTTCCTAC | GAGTTGCTGTTGAAGTCGCA |
| cKit | Mouse | TGGTCAAAGGAAATGCACGA | GCTCCCAGAGGAAAATCCCAT |
| Gata1 | Mouse | TGGGGACCTCAGAACCCTTG | GGCTGCATTTGGGGAAGTG |
| Rps29 | Mouse | GTCTGATCCGCAAATACGGG | AGCCTATGTCCTTCGCGTACT |
| Rps15 | Mouse | GTCATTTACAGGATTCGTGTCCG | CAGGCTTGCCGTAAGTTGC |

**Supplemental table 6**

Primers used in RT-qPCR
