## Supplementary material for "RNA helicase Ddx21 safeguards fetal HSPC expansion by recruiting Kdm5a to epigenetically sustain ribosomal and hematopoietic gene transcription": Figure legends

**Figure 1.** **Ddx21 knockout in fetal hematopoietic cells results in severe hematopoietic defect and embryonic lethality**

(A) Percentage of *Vav1-Cre; Ddx21^f/f^* (*Ddx21*^cKO^) embryos generated from mating pair: *Vav1-Cre; Ddx21^f/+^* (*Ddx21*^Het^) x *Ddx21^f/f^* (*Ddx21*^WT^). N represents the total number of embryos genotyped. (B) Representative gross morphology of whole embryo and fetal liver of *Ddx21*^WT^ and *Ddx21*^cKO^ embryos at E13.5 (n=8) and E18.5 (n=3). (C) Cellularity (n=3-14) of fetal liver of *Ddx21*^WT^, *Ddx21*^Het^ and *Ddx21*^cKO^ embryos. (D) Percentage of dead cells (n=3-14) in fetal liver. (E) Hemoglobin content of whole fetal liver (n=5-8); p-value: WT vs KO. (F) Percentage of Lineage^–^Sca1^+^cKit^+^ (LSK) and Lineage^–^Sca1^–^cKit^+^ (LS^–^K) cells relative to total live fetal liver cells; p-value: WT vs KO. Representative plots of LSK and LS^–^K populations in live Lineage^–^ cells (n=3-8) were shown. (G) LSK, LS^–^K, erythroid and leukocyte populations in total live fetal liver cells (n=5-8). (H) Representative plots of E16.5 erythroid fractions in live Lineage^–^ cells (n=5-8). (I) Representative plots and quantification of E16.5 live Lineage^–^ Ter119^+^ erythroid cells. (J-L) Transplantation of *Ddx21*^WT^ and *Ddx21*^cKO^ fetal liver cells from E13.5 littermate embryos (n=6). (J) Schematic diagram of fetal liver cell transplantation. (K) Peripheral blood (PB) chimerism among CD45^+^ cells at 4 weeks post-transplantation. Representative flow plots of live CD45^+^ cells were shown. (L) Bone marrow (BM) chimerism at 4 weeks post-transplantation. (M-N) Competitive transplantation of *Ddx21*^WT^ and *Ddx21*^cKO^ fetal liver HSCs (live, LSK, CD48^–^, CD150^+^) from E13.5 littermate embryos carrying *mTmG^f/+^* allele (n=6). (M) Schematic diagram of competitive HSC transplantation. (N) PB chimerism among CD45^+^ cells monitored for 12 weeks post-transplantation. (O) BM chimerism at 12 weeks post-transplantation. Representative flow plots of live BM cells were shown. (P) CFU assay of E13.5 FACS-sorted fetal liver Lineage^–^cKit^+^ (LK) cells of *Ddx21*^WT^, *Ddx21*^Het^ and *Ddx21*^cKO^ embryos using MethoCult^TM^ M3434 (n=3-5). Numbers in flow gates represent mean percentage of the population. *p<0.05, **p<0.01, ***p<0.001.

**Figure 2.** **p53 inhibition prevents ROS and p-p53(Ser18)-driven apoptosis and partially restores hematopoietic function**

(A-D) Transcriptomic analysis of *Ddx21*^WT^ and *Ddx21*^cKO^ E13.5 fetal liver LK cells (n=3). (A) Volcano plot showing the differentially expressed genes (DEGs). (B) Top 5 KEGG enrichment in upregulated DEGs (Fold change > 2, p.adj <0.05) in *Ddx21*^cKO^. (C) KEGG p53 signaling pathway GSEA analysis. (D) Expression changes of KEGG p53 signaling pathway genes. (E) Analysis of apoptosis in E13.5 fetal liver cells. Numbers in gates represent mean percentage of total cells (n=3). (F) Flow cytometry analysis of ROS by 2’,7’-dichlorodihydrofluorescein diacetate (H_2_DCF-DA) staining of E16.5 fetal liver erythroid fractions. Geometric mean fluorescence index was reported (n=3-4). (G) Expression of ROS response genes (*Jun*, *Egr1* and *Hif1a*) in E13.5 LK cells (n=3). (H) Immunofluorescence staining of E13.5 live LK cells with p53 and p-p53(Ser18) (n=126-152). (I) Gene expression analysis of *Cdkn1a*, *Mdm2* and *Bax* in cultured E13.5 LK cells after 3 days of Ddx21 knockdown and 15µM Pifithrin-α (PFT-α) incubation (n=3). (J) Colony formation analysis by CFU assay using cultured E13.5 *Ddx21^f/f^* fetal liver LK cells after 3 days of lentivirus-mediated *Ddx21* knockout and p53 knockdown (n=3). Knockdown efficiency was assessed by RT-qPCR analysis of sorted cells. (K) Schematic diagram of human HSPC (live CD34^+^ CD45^+^ CD43^+^) differentiation from hESC. Representative day12 brightfield image overlayed with shRNA-mCherry fluorescence was shown. (L) CFU assay using human HSPC with DDX21 knockdown (shDDX21-1/2) (n=3). (M) Transcriptome analysis by GSEA comparing control (shNC) and DDX21 knockdown (shDDX21) human HSPC (n=3). (N) Flow cytometry analysis of apoptosis in human HSPC after 2 days of drug incubation (n=3). (O) Analysis of hematopoietic capacity by CFU assay using human HSPC with DDX21 and p53 knockdowns (n=3). *p<0.05, **p<0.01, ***p<0.001

**Figure 3.** **Ddx21 loss inhibits protein synthesis and RNA transcription in hematopoietic progenitors**

(A) Enrichment of ribosome biogenesis GO term in mouse HSPC transcriptome by GSEA analysis. (B) Expression profile (by RNA-seq) of the “ribosome biogenesis” geneset (GO:0042254) in the mouse transcriptome. Only significant DEGs (p.adj <0.05) were shown. (C) Venn diagram showing the overlap between mouse and human HSPC DEGs in “ribosome biogenesis” GO term. (D) Heatmap showing all expressed RP genes in mouse HSPC. (E) Quantification of rRNA level in E13.5 LK cells by RT-qPCR (n=6). (F) Tapestaion quantification of rRNA level of E13.5 LK cells (2 x 10^4^) after 3 days of Ddx21 knockdown with or without treatment with 15µM PFT-α (n=3). (G) Enrichment of mitosis and cell cycle related GO terms in mouse HSPC transcriptome. (H) EdU/DAPI-based cell cycle analysis of E13.5 fetal liver live LS^–^K cells 1.5hr after EdU intraperitoneal injection to pregnant mouse. Numbers in the gates represent mean percentage in parental gate (n=3). (I) Enrichment of (downregulated) mitosis and cell cycle related GO terms in human HSPC transcriptome. (J) Analysis of nascent protein synthesis in erythroid fractions by O-propargyl-puromycin (OP-Puro) incorporation assay after 30min of OP-Puro (10µM) incorporation using isolated E13.5 fetal liver cells (n=3). (K) Enrichment of (downregulated) porphyrin-containing compound metabolism and erythrocyte differentiation GO terms in mouse HSPC transcriptome. (L) GSEA analysis on GO term related to “transcription elongation by RNA polymerase II” in mouse HSPC transcriptome. (M) Nascent RNA transcription assay by RT-qPCR on 5-ethynyluridine (EU) incorporated RNA. E13.5 fetal liver LK cells were cultured in 10µM EU for 1hr. Nascent RNA was normalized with total RNA (*Gapdh* RNA before streptavidin bead capture) to calculate relative nascent RNA transcription (n=3). *p<0.05, **p<0.01, ***p<0.001

**Figure 4.** **DDX21 interacts with histone demethylase Kdm5a in an RNA-dependent manner**

(A-C) Identification of DDX21 interaction proteins in K562 cells by proximity-based BioID2 method. (A) Schematic diagram showing the principle of BioID assay. Biotinylated proteins were captured by strepavidin beads and trypsin-digested for mass spectrometry analysis (n=3). (B) Top 5 enriched GO terms of the biotinylated proteins by biological process (BP) and molecular function (MF). (C) Proteins related to histone modifications were enriched in the interactome. (D-G) Co-immunoprecipitation (Co-IP) analysis between DDX21 and KDM5A/HDAC1. Co-IP of Flag-DDX21 with HA-KDM5A (D) and HDAC1 (E) in 293F with or without RNase A treatment. (F) Co-IP of endogenous DDX21 with KDM5A and HDAC1 in K562 cells with or without RNase A treatment. (G) Direct protein pulldown using recombinant proteins of DDX21 with KDM5A or HDAC1. (H) Co-IP between flag-DDX21 mutants and endogenous KDM5A or HDAC1 in 293F. (I) Immunofluorescence staining of endogenous Kdm5a, Hdac1 and Ddx21 proteins in E13.5 fetal liver LK cells with or without (mock) RNase A treatment. Quantification of the relative fluorescence unit (RFU) was shown (n=78-108). Scale bar: 5µm. (J) Proximity ligation assay (PLA) staining for Ddx21-Kdm5a and Ddx21-Hdac1 interactions. LK cells were either mock or RNase A-treated before staining. Quantification of PLA RFU was shown (n=65-95). Scale bar: 5µm. *p<0.05, **p<0.01, ***p<0.001

**Figure 5.** **Ddx21 and Kdm5a co-occupy active promoters to preserve a transcriptionally active state**

(A-H) Epigenetic profiling and comparative analysis of H3K4me3, H3K4me1, H3K27me3, H3K27Ac, and Kdm5a binding by CUT&Tag-seq. ATAC-seq was also performed in E13.5 fetal liver LK cells (n=2, biological replicates). Signals of the same target were normalized, scaled to per-million reads and merged between replicates. Promoters were defined as +/-1500bp of transcription start site (TSS). (A) Pearson correlation of all signals (bigWig) between histone marks, Ddx21 and Kdm5a in *Ddx21*^WT^ cells. (B) Profile plot of CUT&Tag and ATAC signals relative to Ddx21 binding in *Ddx21*^WT^ cells. (C) Peak number (nPeak) and distribution of H3K4me3, Ddx21 and Kdm5a binding peaks in *Ddx21*^WT^ cells. (D) Pearson correlation of log_2_(normalized counts+1) in all promoters of *Ddx21*^WT^ cells between Kdm5a and Ddx21, H3K4me3, and RNA. (E) Profile plots of H3K4me3, H3K4me1, H3K27Ac, H3K27me3, ATAC-seq, and Kdm5a at TSS+/-3kb, comparing *Ddx21*^WT^ and *Ddx21*^cKO^ cells. (F) Profile plots and heatmaps of active (H3K4me3+, H3K27me3-, RNA+), poised (H3K4me3+, H3K27me3-, RNA-), bivalent (H3K4me3+, H3K27me3+) and repressive (H3K4me3-, H3K27me3+) promoters in *Ddx21*^WT^ and *Ddx21*^cKO^ cells. Differential gene expression (RNA) in these clusters was also shown. (G) Clusters of Ddx21-bound promoters based on Kdm5a-binding changes (*Ddx21*^cKO^ vs *Ddx21*^WT^, Log_2_FC > 0.2 or < -0.2). Heatmaps of Kdm5a occupancy, H3K4me3 and RNA expression were shown. Violin plots of the respective clusters were shown; values represent mean of Log_2_FC in each cluster. (H) GO enrichment comparison (q<0.05) of the genes from the 3 clusters. *p<0.05, **p<0.01, ***p<0.001

**Figure 6.** **Dual inhibition of Kdm5a and p53 restores rRNA level and translational capacity in Ddx21-deficient hematopoietic progenitors**

(A) The genomic landscape of Ddx21 and Kdm5a occupancy, and the related histone marks in ribosomal DNA (rDNA) tandem repeat of E13.5 *Ddx21*^WT^ and *Ddx21*^cKO^ fetal liver LK cells (n=2). IGS: intergenic spacer; SpPr: spacer promoter; 47S-Pr: 47S rRNA promoter; ETS: external transcribed spacer; ITS: internal transcribed spacer; TSS: transcription start site; TES transcription end site. (B-D) Flow cytometry analyses of nascent protein synthesis, apoptosis, and cell cycle of cultured E13.5 fetal liver LK cells after 3 days of Ddx21 knockdown (shDdx21) and incubated with 15µM PFT-α / 5-10µM CPI-455 / DMSO (n=3). (B) Median fluorescence of OP-Puro in live shRNA^+^ cells after 30min of OP-Puro incorporation. (C) Percentage of dead or pre-apoptotic cells (determined by Apotracker and fixable viability dye staining). (D) EdU/DAPI-based cell cycle analysis of live shRNA^+^ cells after 1hr of EdU incorporation. (E) CFU assay of cultured E13.5 *Ddx21^f/f^* fetal liver LK cells after 3 days of Cre-mediated knockout of Ddx21 and shRNA-mediated knockdowns of Kdm5a and p53. Live shRNA^+^ cells were FACS-sorted for CFU plating (n=3). (F) Flow cytometry analysis of E13.5 fetal liver after 3mg/kg CPI-455 and 2.2mg/kg PFT-α injections at E11.5 and E12.5. Representative flow plots of live lineage^–^ cells were shown. Numbers in red wer mean percentages of the parent gate. Quantifications (relative) of LS^–^K and LSK cells were shown (n=4-6). *p<0.05, **p<0.01, ***p<0.001
